# One digit too many: brain rhythms and autonomic responses at the limits of working memory capacity

**DOI:** 10.64898/2026.11.01.763481

**Authors:** Alexander I. Kotyusov, Dauren A. Kasanov, Alexandra I. Zaitseva, Boris Kotchoubey, Yuri G. Pavlov

## Abstract

Physiological measures vary with working memory load, but whether they identify when demands exceed a person’s working memory capacity remains unclear. We assessed neural and autonomic responses to exceeding digit span by one digit. During an adaptive digit span task, we recorded EEG, pupillometry, and photoplethysmography from 123 participants. We compared each participant’s digit span (approximately 70% correct full-sequence recall) with sequences one digit longer (span+1), operationally defined as overload. Full-sequence accuracy decreased from 74.1% to 51.5%, and subjective mental demand increased. Baseline-normalized pulse wave amplitude (PWA) was lower in span+1 from the first digit through terminal retention period. PWA was lower on correctly recalled than erroneous span+1 trials. Phasic pupil responses increased across serial positions but were smaller in span+1 at late span-matched positions; responses did not differ between conditions at their final digits. Non-baseline-corrected pupil diameter was larger in span+1 before sequence onset and during terminal retention, whereas non-baseline-corrected PWA did not differ. Frontal midline theta increased with serial position and was greater during terminal retention in span+1. Alpha and beta power decreased with serial position, and differed between conditions only at late positions. Terminal beta suppression was greater on correctly recalled span+1 trials. Thus, exceeding calibrated span impaired recall without producing a common physiological breakpoint. The findings suggest that reaching this limit does not immediately lead to resource exhaustion or disengagement. Physiological responses showed shifts in timing consistent with adjustments in preparation for recall of the longer sequence, along with increased effort before and during longer sequences.

## Introduction

Working memory allows us to temporarily store a limited amount of information in mind for later use (Cowan, 2001, 2010). This storage capacity limit varies across individuals, sensory modality, and type of task, but healthy young adults typically recall about six or seven digits in a standard digit span task (Gignac & Weiss, 2015). When digit sequence length exceeds an individual’s digit span, performance does not immediately collapse: accuracy often remains well above floor, although recall of each item in the longer sequence is impaired (e.g., Kosachenko et al., 2023). This pattern suggests that working memory overload is best understood as a state in which people remain engaged while the processes supporting working memory are placed under increasing strain (Oberauer, 2019; Oberauer et al., 2016; Oberauer & Lewandowsky, 2011). This boundary state – when task demands exceed this capacity by the smallest measurable amount, such as one additional digit in a digit span task – remains poorly characterized in both behavioral and psychophysiological research.

To understand what happens at the boundary of capacity, behavioral measures are frequently insufficient because measures such as accuracy and reaction time only describe the end result of the task performance rather than the processes that produced it. They, therefore, need to be complemented by physiological measures that follow neural and autonomic activity as the sequence unfolds. Pupil diameter is among the most established indices of cognitive effort: it dilates as more information is held in memory (Kahneman & Beatty, 1966), and later studies suggested that this response can plateau or reverse when task demands exceed available resources (Cabestrero et al., 2009; Granholm et al., 1996; Kosachenko et al., 2023). Pulse wave amplitude, derived from peripheral photoplethysmography, provides a complementary autonomic index decreasing with sympathetically driven vasoconstriction during effortful cognition (Allen, 2007; Charkoudian, 2003) and shown to track cognitive load in digit span task specifically (Pavlov et al., 2023). EEG provides temporally precise information about cortical processes involved in serial encoding and retention. Frontal midline theta activity typically increases with working memory and cognitive control demands and is associated with executive control and conflict monitoring (Cavanagh & Frank, 2014; Roux & Uhlhaas, 2014). Posterior alpha activity is commonly associated with attentional selection, inhibition of task-irrelevant stimulus processing, and controlled access to stored information (Jensen, 2023; Klimesch, 2012). Sensorimotor beta suppression is associated with motor preparation (Boettcher et al., 2021; Kilavik et al., 2013; Schneider et al., 2017). Whether exceeding memory span by a small margin increases, stabilizes, or attenuates these signals is not known.

This question is difficult to answer in fixed-load designs because working memory capacity varies substantially between individuals, including in digit span (Gignac & Weiss, 2015; Unsworth & Engle, 2007). In a serial recall task such as digit span, a fixed sequence length may fall within one participant’s reliably recalled range, exceed another participant’s span by one item, and exceed a third participant’s span by several items. A group level average can therefore combine qualitatively different states: manageable demand, the narrow transition at capacity, deeper overload, and, at sufficiently high difficulty, possible disengagement. Our own earlier work using fixed sequence lengths revealed a clear multimodal physiological signature of overload (Kosachenko et al., 2023), but precisely because the degree of exceedance varied across individuals, it could not isolate what happens at the boundary of capacity itself, when span is exceeded by the smallest possible margin. Defining overload relative to each participant’s own span reduces variation in relative task difficulty, providing a more precise test of whether a physiological overload signature is present at the very edge of capacity. Beyond its theoretical value, this individualized approach may provide a foundation for personalized cognitive load monitoring.

The present study tested how a slight, individually calibrated exceedance of working memory capacity changes behavioral performance, subjective demand, and physiological dynamics during serial digit encoding and retention. Participants completed an adaptive digit span task that established two adjacent sequence lengths, designated span and span+1. We use “overload” operationally to denote the span+1 condition: a sequence one digit longer than the participant’s adaptively selected reference span. We expected lower full-sequence and position-wise recall accuracy and higher subjective mental demand at span+1. Observing this pattern would serve as a manipulation check. The physiological analyses were primarily exploratory, asking whether the clear multimodal data signature seen under overload persists at the very edge of capacity, or whether minimal, individually calibrated overload reveals a different picture. For each measure, we asked whether activity would increase with the added demand, plateau at a functional limit, or decline, with interpretations varying depending on the specific physiological measure.

## Methods

### Design

The study used a within-subject design and was conducted across two visits. During the first visit, participants completed a cognitive task battery, including a simple digit span task that provided an initial estimate of each participant’s individual span (see Kotyusov et al., 2023, for a detailed description). The second visit included an adaptive working memory task with simultaneous physiological recordings of electroencephalography (EEG), pupillometry, electrocardiography (ECG; not analyzed here), and photoplethysmography (PPG).

### Participants

In total, 137 participants (111 females) were recruited for this study at Ural Federal University. Most participants were university students. All participants were native Russian speakers, had normal or corrected-to-normal vision, and reported no history of neurological or psychiatric disorders. The study protocol was approved by the ethics committee of Ural Federal University, and all participants provided written informed consent.

Twelve participants did not return for the second visit. In addition, one participant was excluded due to intentional random responding, and one participant was excluded because of technical problems. Thus, fourteen participants were excluded from all analyses, leaving a final sample of 123 participants (100 females), aged 18 to 32 years (M = 20.49, SD = 2.27). Some participants were excluded from specific modalities, as described below.

Handedness was assessed using the Annett Handedness Scale (Annett, 1970). Four participants were ambidextrous, 10 were left-handed, and 109 were right-handed. Ocular dominance was determined using three tests: the Porta-Rosenbach alignment test, an aiming test in which participants made a finger-gun gesture with both hands and aimed at a self-selected target while closing one eye at a time (Miles, 1930), and the hole-in-the-card test (Fink, 1938). Forty-five participants had left-eye dominance, and 78 had right-eye dominance.

Three participants were excluded from the behavioral analyses because of technical errors in data saving. Thus, 120 participants were included in the behavioral analyses. Twenty of these participants had technical problems with the NASA-TLX subjective mental demand scale. Therefore, the final sample size for the NASA-TLX analysis was 100 participants.

For all physiological analyses, but not the behavioral analyses, one participant was excluded because of hardware failure that left too few trials for analysis. The final sample size for the EEG analysis was 122 participants, including 119 participants with both behavioral and EEG data and three participants without behavioral data. For these three participants, individual load level and span length were extracted from the event markers in EEG data.

Because of a technical error during recording, PPG data were missing for three participants. Since PPG data were recorded as part of the EEG recordings, the final sample size for this modality was 119 participants.

Two participants were excluded from the pupillometry analysis due to missing data. Participants with fewer than 20 trials in either the span or span+1 (overload) condition were also excluded. Applying these exclusion criteria left 109 participants in the pupillometry sample.

### Task and procedure

In each trial, participants were presented with a sequence of digits to be remembered and recalled, similar to the simple digit span (see Figure 1). The trial began with an exclamation mark for 0.5 s, followed by a fixation cross (Thaler et al., 2013) (0.8 cm in diameter) for 2 s, and then digits were presented for 0.5 s with SOA of 2 s alternated with the fixation cross. After the presentation of the last digit, the retention interval continued for 3.5 s. Participants were then required to type the sequence in the correct order, with no time limit for responding. Participants could correct their input, but were required to enter the full sequence. A successful trial was defined as recalling all digits in the correct order.

**Figure 1.**
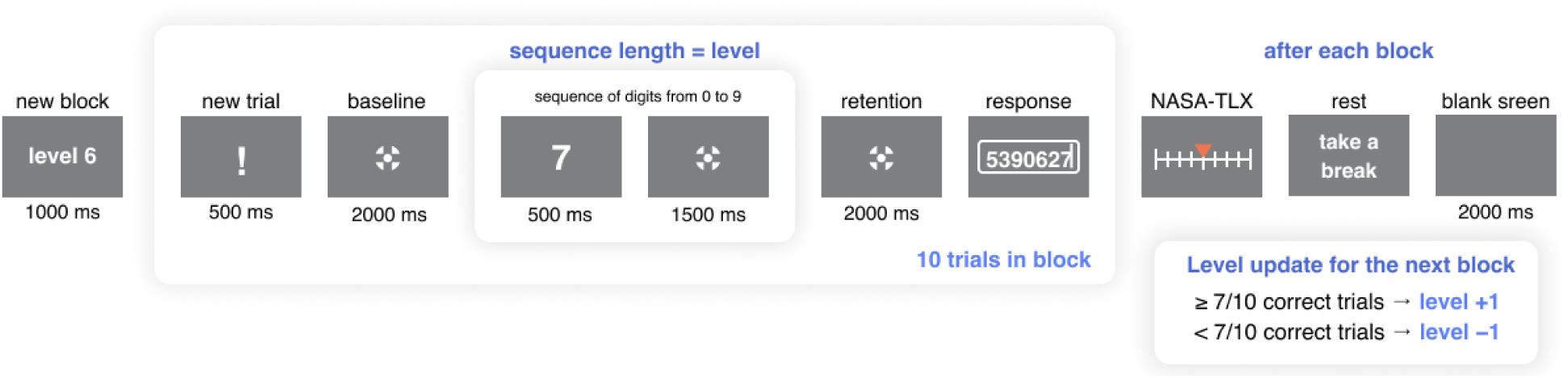
Adaptive digit span task and trial timeline. Before each 10-trial block, participants were informed of the sequence length, presented as a task “level.” Each trial began with an exclamation mark (0.5 s) and a fixation baseline (2 s). Digits from 0 to 9 were then presented for 0.5 s each, separated by 1.5-s fixation intervals. Following the final digit and its fixation interval, an additional 2-s retention period produced a total terminal retention interval of 3.5 s before the response screen. Participants typed the complete sequence in serial order without a response time limit. After each block, they rated mental demand on the 0-20 NASA-TLX scale and could take a self-paced break. Sequence length increased by one after at least 7 of 10 completely correct trials and decreased by one otherwise. The diagram is not to scale.

**Figure 2.**
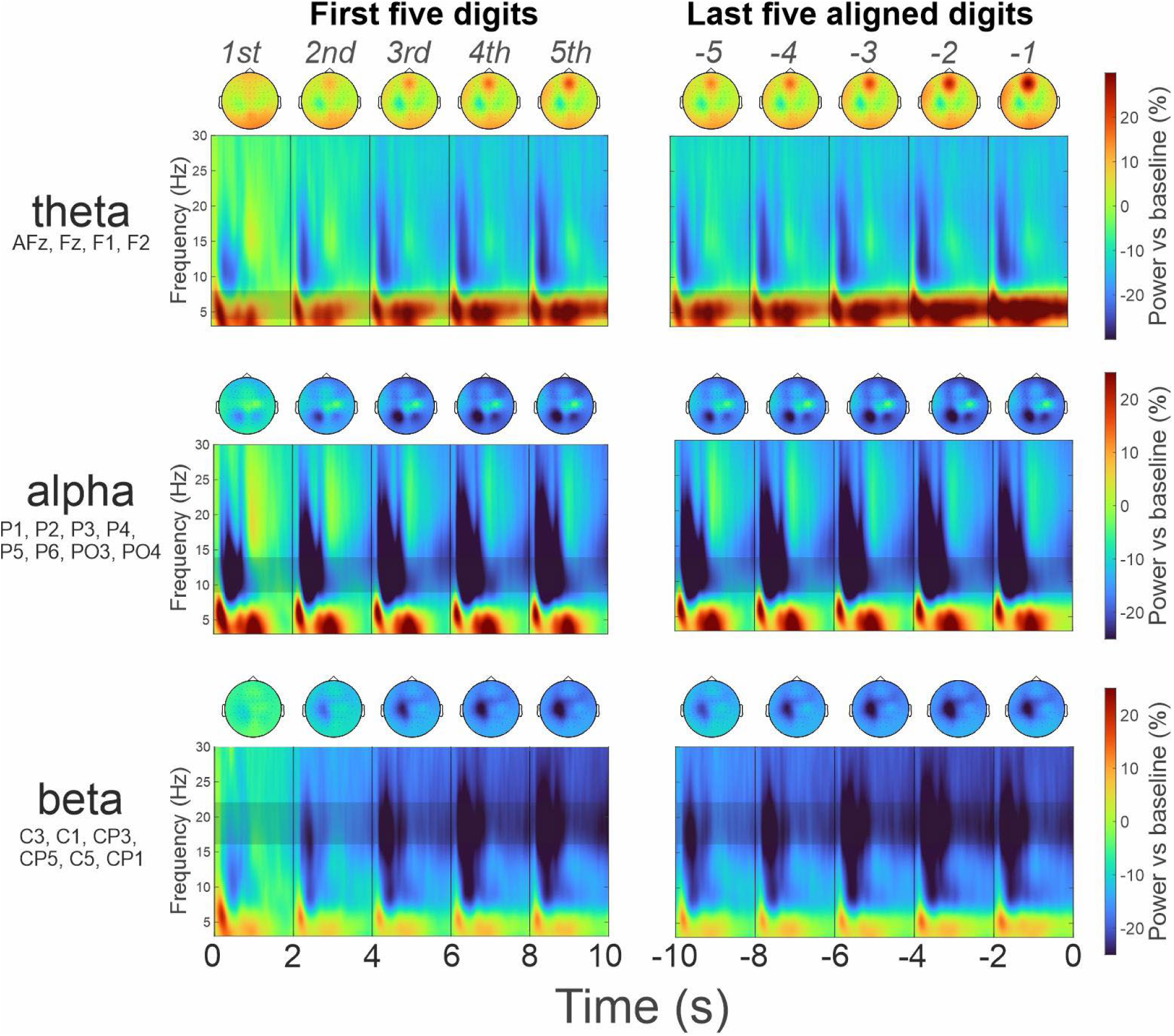
Grand-average EEG time-frequency dynamics (flattened average) used to define the frequency bands and spatial regions of interest. Time-frequency power and corresponding scalp distributions are shown for frontal-midline theta (4-8 Hz; AFz, Fz, F1, and F2), posterior alpha (9-14 Hz; P1, P2, P3, P4, P5, P6, PO3, and PO4), and left centroparietal beta (16-22 Hz; C3, C1, CP3, CP5, C5, and CP1). The left column shows responses to the first five digits; the right column shows the five late, span-aligned positions used in the serial-position analyses (−5 to −1). Under the primary span/span+1 convention, position −1 is the final span digit and position 0 would be the additional span+1 digit. Power is expressed as percentage change from the 1-s presequence baseline. Scalp maps summarize band-limited activity during the corresponding 2-s digit intervals. The flattened averages include all valid trials from all observed sequence lengths, collapsed across conditions and participants (N = 122)

Participants completed a simple digit span task during the first visit. In the adaptive digit span task administered at the second visit, their starting span length was defined as their maximum correctly recalled span length from the first visit minus one digit.

Trials Trials were presented in blocks of 10 sequences of the same length. Participants were informed of the sequence length before each block (in a gamified way presented as levels - e.g., “level 6” would simply mean there will be 10 trials with 6 digit sequences). Sequence length increased by one digit after a block with at least seven fully correct responses and decreased by one digit otherwise. The task ended once participants had completed at least six blocks (60 trials) at each of two adjacent sequence lengths. The shorter length was designated span and the longer span+1 (overload). All trials at these two lengths were included in the analyses, regardless of the accuracy achieved in individual blocks.

Operationalizing cognitive overload requires a decision about the accuracy threshold used to define an individual’s digit span. In the primary analyses, the digit span was defined as the sequence length at which participants fully recalled at least 7 of 10 sequences (≥70% accuracy). This criterion is more stringent than the conventional WAIS-IV Digit Span termination rule, under which testing at a given list length ends after two errors and one correct response out of two trials is sufficient (50% accuracy). The 70% criterion was selected to provide a more stable estimate of reliably manageable span (Sebastián & Mediavilla, 2015; Woods et al., 2011).

Because the task was adaptive, some participants completed more than the minimum of 120 trials. This, together with self-paced breaks between blocks, contributed to variability in task duration. On average, 66 trials from span blocks and 66 trials from span+1 blocks were included in the analysis.

After each block, participants answered one item from the NASA-TLX mental demand scale (Hart & Staveland, 1988): “How mentally demanding was the task?” The visual analog scale ranged from 0 to 20.

The dependent variables were full-sequence accuracy and partial recall accuracy. Full sequence accuracy was defined as whether the entire sequence was recalled in the correct order. Partial recall accuracy was calculated as the number of digits recalled in the correct serial position, independently of sequence length. For example, if the presented sequence was 416058972 and the response was 40615897, the partial score was 6.

The viewing distance was 80 cm. Room luminance was set to 380 lux. The horizontal font size was 1.1 visual degrees. The screen measured 23.6 inches diagonally. Throughout the task, the background color was gray (RGB: 128, 128, 128), and the font color was white (RGB: 255, 255, 255). The task was presented using PsychoPy software (Peirce et al., 2019).

### Electroencephalography

EEG was recorded using a 64-channel active electrode system (actiCHamp, Brain Products). Electrodes were placed according to the extended 10-20 system, with Fpz used as the ground electrode. The FCz channel was used as the online reference in most participants. However, in 20 participants, Fp2 was used as the reference because of human error. The reference channel was reconstructed before average re-referencing, so the online reference does not affect the analysed data, Electrode impedance was kept below 25 kOhm. The sampling rate was 1000 Hz.

EEGLAB (Delorme & Makeig, 2004) was used for data preprocessing. Each recording was filtered with a 1 Hz high-pass filter and a 45 Hz low-pass filter (with default EEGLAB settings). Visually identified bad channels were spherically interpolated. The data were then downsampled to 250 Hz, re-referenced to the average reference, and subjected to independent component analysis using the AMICA algorithm (Palmer et al., 2012). Components related to eye movements and blinks were removed. In addition, components mapped to a single electrode and clearly distinguishable from EEG signals were subtracted from the data. The data were epoched from −1500 to 5000 ms relative to digit presentation to avoid edge artifacts. Epochs that still contained artifacts were visually identified and discarded.

Time-frequency analysis was performed separately for each participant and condition on the preprocessed single-trial data, from 1 to 45 Hz in 1-Hz steps, using Morlet wavelets whose number of cycles increased logarithmically from 3 at 1 Hz to 12 at 45 Hz; the window was shifted in 20-ms steps. Spectral power was baseline-normalized by computing the percent change in power relative to the −1000 to 0 ms interval, which corresponded to the last second of the baseline fixation period before presentation of the first digit in a sequence. Time-frequency analysis was performed using the FieldTrip toolbox (Oostenveld et al., 2011).

For EEG analyses, three frequency bands and regions of interest (ROIs) were selected: frontal midline theta (4-8 Hz; AFz, Fz, F1, and F2), posterior alpha (9-14 Hz; P1, P2, P3, P4, P5, P6, PO3, and PO4), and beta (16-22 Hz; C3, C1, CP3, CP5, C5, and CP1). The centro-parietal cluster was selected because this region showed the most pronounced beta suppression after averaging across all participants and conditions. A similar approach was used for frontal midline theta. The alpha ROI covered a large part of the occipito-parietal cortex, where alpha suppression and rebound during the encoding and delay periods was visually prominent on the flattened average (Bowman, Brooks, Hajilou, Zoumpoulaki, & Litvak, 2020).

### Pupillometry

Pupillometry was recorded using a Pupil Labs wearable eye tracker (Pupil Labs GmbH, Germany) at a sampling rate of 120 Hz. A one-point calibration was performed before each recording. During preprocessing, data from the dominant eye were selected. Data points identified by the Pupil Player algorithm as having a confidence score below 80% were replaced with missing values for later interpolation. The data were then segmented into epochs beginning 2 s before presentation of the first digit in the sequence and ending at the onset of the response window.

Further processing was conducted using the Gazer package in R (Geller et al., 2020). Blinks detected by the Pupil Labs algorithm were extended by 100 ms before and 100 ms after each blink and replaced with missing values. Pupil values exceeding 16 median absolute deviations were also rejected. Trials containing more than 50% missing values were removed from the analysis. Participants with fewer than twenty remaining trials in any condition were excluded from the analysis. Missing values in the remaining trials were linearly interpolated, and the data were smoothed using a five-point moving average. Pupil size was baseline-normalized by subtracting the mean absolute value during the 1 s interval before presentation of the first digit in the sequence. The data were then downsampled to 10 Hz. For the tonic analysis of raw pupil diameter (see Statistics), the non-baseline-corrected diameter was retained after the same artifact rejection, interpolation, smoothing, and downsampling steps.

### Photoplethysmography

Finger photoplethysmography was recorded from the left middle finger using the auxiliary inputs of the actiCHamp amplifier (Brain Products). Preprocessing of PPG included the following steps. First, PPG data were filtered between 0.5 and 100 Hz, and a 50 Hz notch filter was applied using the eegfiltnew function in EEGLAB. The data were then downsampled to 250 Hz. Next, HRVTool v.1.07 (Vollmer, 2019) was used to identify troughs and peaks in the PPG signal. Each trough and peak was visually inspected, and incorrectly detected values were rejected from further analysis. Interbeat intervals shorter than 0.3 s or longer than 2.4 s, as well as pulse wave amplitude (PWA) values more than 3 SDs above or below the mean, were also rejected.

PWA was defined as the amplitude difference between consecutive troughs and systolic peaks, starting from the trough. The recording was then segmented into trials beginning 3.5 s before presentation of the first digit and ending 1.5 s after response window onset. Within these epochs, PWA series were linearly interpolated in 0.5 s bins to obtain equally spaced PWA time series. Trials with 10 or more rejected values at the previous steps were not interpolated and removed from the analysis.

For the time-locked analyses, after averaging across trials, the 1 s interval before presentation of the first digit was used for baseline normalization. Baselines were calculated separately for each participant and sequence length. Relative baseline normalization (percent change) was applied to PWA values. For the tonic analysis of raw PWA (see Statistics), non-normalized PWA values were retained.

### Statistics

For the statistical analyses of EEG, PPG, and pupillometry data, relative spectral power in three frequency bands, pulse wave amplitude (PWA), and pupil size were averaged in 2 s intervals corresponding to the encoding and maintenance of individual digits. For the analyses exploring the effect of overload on the final digit in the sequence, values were averaged over the 4 s interval before the recall probe.

Primary analyses were conducted using repeated measures ANOVAs. The effect of the additional final item in the overload sequence was assessed in an ANOVA that included only the final digit from the span and overload sequences.

Two more ANOVAs were conducted to study changes across serial positions. These analyses included the factors Serial Position (five levels), and Condition (span vs overload). For the analysis of the first five digits, physiological activity and accuracy during processing of the first five digits were included. For the analysis of the last five digits, we compared span and overload trials at the same item positions counted from the end of the span sequence length. In this analysis, the final item from the overload sequence was not included in the analysis (although visualized in figures) and the digits were taken on positions corresponding to the last five positions of span condition. Such an approach allowed us to assess the effect of the sequence context. Because two participants had relatively low digit spans of 3 or 4 digits, they were excluded from the serial position x condition analyses.

Because per-condition baseline normalization removes any block-level difference between conditions, we compared the raw, non-baseline-corrected pupil diameter and PWA between span and span+1 in two intervals: the 2-s pre-sequence period (−2 to 0 s) and the terminal encoding and retention interval (the 4 s from the onset of the final digit to the recall probe). Each participant’s raw signal was z-scored using the mean and SD of all samples from all of that participant’s span and span+1 trials. Interval means were computed for each trial, averaged per participant and condition, and compared with paired t-tests.

To test whether physiological activity predicted recall success under overload, span+1 trials that were recalled fully correctly were compared with span+1 trials containing at least one error, in the terminal encoding and retention interval and across the first and last five digits of the span+1 sequence, with recall correctness (correct vs. error) as the within-subject factor.

Greenhouse-Geisser corrected degrees of freedom were reported when the assumption of sphericity was violated. Post hoc pairwise comparisons with paired t-tests were conducted where appropriate. All statistical analyses were performed in R, version 4.4.2.

## Results

### Behavior

#### Accuracy

The number of fully recalled sequences decreased in the span+1 (overload) in comparison with the span condition (F(1, 119) = 351.54, p < 0.001, *η*_p_^2^ = 0.75), as well as percent accuracy (F(1, 119) = 217.42, p < 0.001, *η*_p_^2^ = 0.65). As can be seen in Table 1, the mean (±SD) proportion of fully recalled sequences decreased from 74.1% ± 8.9 in the span condition to 51.5% ± 9.7 in the overload condition, similarly to the partial recall: 90.5% ± 4.3 vs 82% ± 5.5 in the span and overload conditions, respectively. The absolute number of correctly recalled digits was higher in the overload condition than in the span condition (6.38 ± 1.28 vs 6.12 ± 1.25, F(1, 119) = 38.64, p < 0.001, *η*_p_^2^ = 0.25).

**Table 1.** Behavioral performance and subjective mental demand ratings in the span and span+1 conditions.

| Condition | Mental demand, M (SD) | Partial-recall accuracy, %, M (SD) | Digits correct, n, M (SD) | Median sequence length | Correct sequences, %, M (SD) |
|---|---|---|---|---|---|
| Span | 10.39 (3.19) | 90.5 (4.32) | 6.12 (1.25) | 6.5 | 74.1 (8.94) |
| Span+1 (overload) | 12.49 (2.88) | 82.0 (5.47) | 6.38 (1.28) | 7.5 | 51.5 (9.65) |

To examine the dynamics of digit encoding and retention between contexts of span and overload conditions, we analyzed recall accuracy for the first five digits as a function of serial position (Figure 3). Because only two participants had a span level below five digits, they were excluded from this analysis. We conducted a 5 × 2 ANOVA with the factors Serial Position and Condition (span vs. overload). There was a significant effect of Condition (F(1, 117) = 123.14, p < 0.001, *η*_p_^2^ = 0.51), with poorer performance in the overload condition. Accuracy also decreased with serial position (F(2.84, 331.79) = 108.25, p < 0.001, *η*_p_^2^ = 0.48), and the gap between overload and span condition increased with serial position (F(3.38, 395.19) = 14.85, p < 0.001, *η*_p_^2^ = 0.11). Pairwise comparisons confirmed a significant decrease in recall accuracy across subsequent digits, except for the difference between the second and third digits in the overload condition.

**Figure 3.**
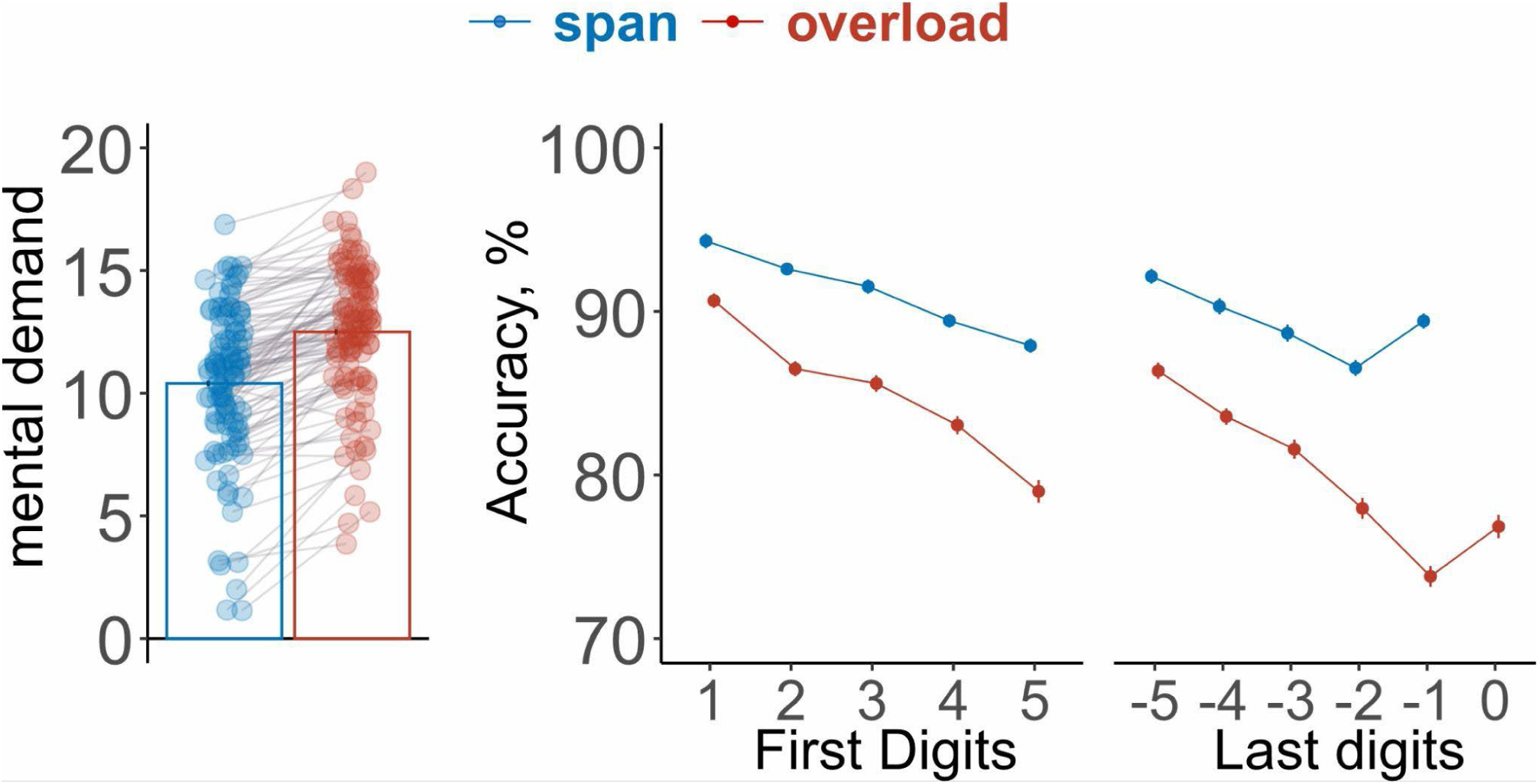
Subjective mental demand and position-wise recall accuracy in individual span and span+1 overload conditions. Blue denotes span and red denotes span+1. The left panel shows NASA-TLX mental demand ratings on a 0-20 scale (N = 100); translucent points are individual participants, gray lines connect repeated measurements, and outlined bars show condition means. The accuracy panels show the mean percentage of digits recalled in their correct serial positions. The first analysis includes positions 1-5 from sequence onset. In the late-position analysis, positions −5 to −1 are the final five positions of the span sequence; position 0 is the additional final digit presented only in span+1. Accuracy declined with serial position and was lower in span+1, with the condition difference increasing toward the end of the sequence. Error bars show SEM.

Recall accuracy for the last five digits was analyzed in the same way (Figure 3). The results largely mirrored those obtained for the first five digits, with lower accuracy in the overload condition (Condition main effect: F(1, 117) = 174.17, p < 0.001, *η*_p_^2^ = 0.60). Accuracy also decreased across serial positions (Serial position: F(3.54, 414.18) = 88.19, p < 0.001, *η*_p_^2^ = 0.43). The significant Serial Position × Condition interaction reflected both a steeper decrease in the overload condition and a prominent recency effect in the span condition (F(3.67, 429.35) = 58.92, p < 0.001, *η*_p_^2^ = 0.33). In the span condition, pairwise comparisons showed that accuracy decreased for digits closer to the end of the sequence, except for the final digit, which was recalled more accurately than the preceding digit. In the overload condition, all serial positions differed significantly, showing a strong downward trend. A separate analysis of the overload sequence over the last two items including the final item in the overload condition confirmed the presence of a recency effect (t(117) = 4.132, p < .001, d = 0.382). This effect was similar to the recency effect observed in the span condition (t(117) = 6.417, p < .001, d = 0.593).

#### Subjective task difficulty

To confirm that the task was perceived as more challenging in the overload condition, we compared NASA-TLX mental demand ratings between the overload and span conditions. The overload condition was rated as subjectively more difficult than the span condition (F(1, 99) = 164.68, p < 0.001, *η*_p_^2^ = 0.62; see Table 1 for descriptive statistics).

### Pupillometry

#### Tonic response

##### Span vs overload: retention interval

The analysis of average pupil response during the retention period after presentation of the final digit revealed no significant difference between the span and overload conditions (F(1, 108) = 0.18, p = .676, *η*_p_^2^ < .01; Figure 4a: right panel).

**Figure 4.**
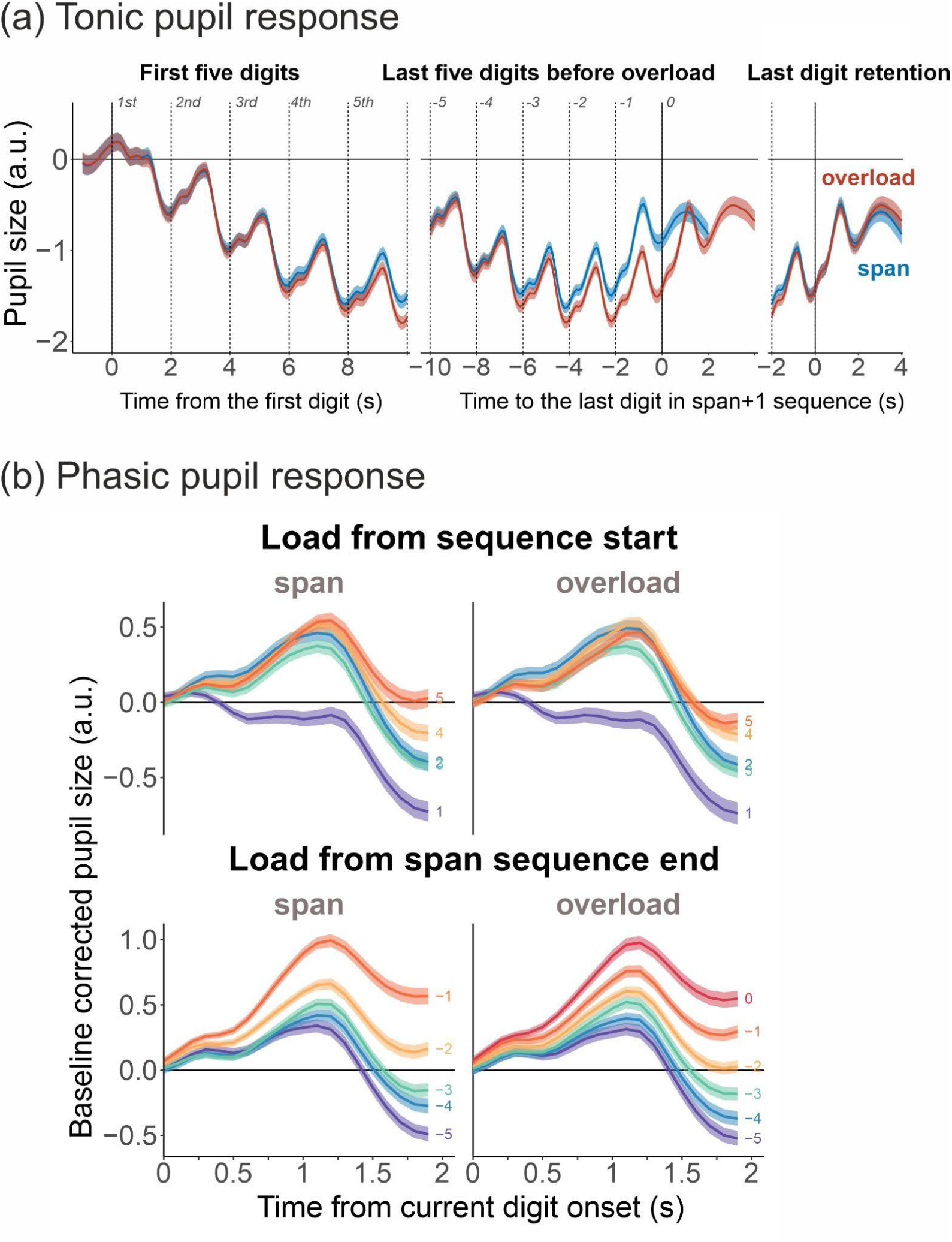
Tonic and digit-evoked pupil responses in individual span and span+1 overload condition. (a) Tonic pupil size, expressed in arbitrary units after subtraction of the 1-s presequence baseline. Traces are shown from the first digit, across the late span-aligned positions, and around the final digit of each condition. Blue denotes span and red denotes span+1; time 0 in the right panel is the onset of each condition’s final digit. (b) Phasic pupil responses obtained by subtracting mean pupil size during the 200 ms immediately preceding each digit. The upper panels show responses to positions 1-5 from sequence onset. The lower panels show the late positions aligned to the span sequence end: −1 is the final span digit and 0 is the additional final span+1 digit. Line colors identify serial position. Phasic responses generally increased with serial position, but were smaller in span+1 at the latest matched positions and did not differ between conditions at their respective final digits. Shaded bands show SEM.

##### First and last five digits

For the first five digits in the sequence, pupil size decreased as working memory load increased (Serial position: F(1.41, 149.57) = 108.67, p < .001, *η*_p_^2^ = .51). There was no significant effect of Condition (span vs overload: F(1, 106) = 1.54, p = .218, *η*_p_^2^ = .01) or Condition × Serial position interaction (F(1.85, 196.45) = 2.88, p = .062, *η*_p_^2^ = .03).

For the last five digits, the main effect of Condition indicated larger pupil size in the span condition than in the overload condition (F(1, 106) = 14.78, p < .001, *η*_p_^2^ = .12). Pupil size decreased until approximately the third digit before the end of the sequence, then plateaued, and increased again at the final digit, as indicated by the main effect of Serial Position (F(1.24, 131.11) = 28.07, p < .001, *η*_p_^2^ = .21).

The dynamics of pupil size changes differed between conditions (Serial position × Condition interaction: F(2.10, 222.79) = 26.97, p < .001, *η*_p_^2^ = .20). Post-hoc t-tests indicated that the difference between the span and overload conditions was significant for the final digit of the span sequence length (t(106) = 2.376, p = .019, d = 0.23), the second-to-last digit (t(106) = 3.610, p = .001, d = 0.35), and the third-to-last digit (t(106) = 7.159, p < .001, d = 0.69). The difference was not significant for the fourth-to-last digit (t(106) = 1.384, p = .169, d = 0.13) or the fifth-to-last digit (t(106) = 0.688, p = .493, d = 0.07). Thus, the dissociation between the overload and span conditions started around the third digit before the end of the sequence.

### Phasic response

Because tonic pupil size unexpectedly showed an overall downward trajectory during visual digit presentation, we conducted a complementary phasic response analysis. In this analysis, pupil responses were time-locked to the onset of each digit and baseline-corrected by subtracting the mean pupil size during the 200 ms preceding digit presentation. This analysis was intended to isolate digit evoked pupil changes from the tonic response trajectory (see Figure 4B).

#### Span vs overload: maximal load

The phasic pupil response did not differ between the final digit in the span condition and the additional digit in the overload condition, (F(1, 108) = 0.07, p = .795, *η*_p_^2^ < .01). Thus, consistent with the tonic pupil analysis of the final retention period, pupil responses converged at the end of the sequence.

#### First and last five digits

For the first five digits, phasic pupil responses increased significantly across serial positions (F(3.14, 332.76) = 22.58, p < .001, *η*_p_^2^ = .18). However, there was no main effect of Condition, (F(1, 106) = 0.92, p = .341, *η*_p_^2^ < .01), and no Condition × Serial Position interaction (F(4, 424) = 1.45, p = .217, *η*_p_^2^ = .01), suggesting a similar trend across conditions. A linear regression analysis confirmed a positive linear increase in phasic pupil responses across the first five digits in both the span (t = 6.26, p < .001) and overload conditions (t = 4.21, p < .001). The linear slopes did not differ significantly between conditions (t = 1.46, p = .145).

The last five digits analysis showed a different pattern. Phasic pupil responses were on average larger in the span than in the overload condition, (F(1, 106) = 26.64, p < .001, *η*_p_^2^ = .20), and generally increased across serial positions, (F(3.48, 369.22) = 46.36, p < .001, *η*_p_^2^ = .30). The Condition × Serial Position interaction was significant (F(3.60, 381.49) = 6.27, p < .001, *η*_p_^2^ = .06) with pairwise comparisons indicating larger responses in the span condition at the two latest matched positions in the analysis (p = .029, d = 0.22, and p < .001, d = 0.56), but not in the other positions. The ordinary linear models also indicated an overall positive linear trend across the last five digits in both the span (t = 3.06, p = .002) and overload conditions (t = 2.90, p = .004). The slopes did not differ between conditions (t = 0.11, p = .914).

Thus, overload was not associated with a uniform reduction or increase in phasic pupil responses. The condition differences were temporally localized with responses being identical for the early analyzed items, differed during presentation of the late items, and converged again at the final item.

### Pulse wave amplitude

#### Span vs overload: retention interval

We found a significant effect of overload, with lower PWA in the overload condition (F(1, 118) = 21.89, p < .001, *η*_p_^2^ = .16; Figure 5, right panel)

**Figure 5.**
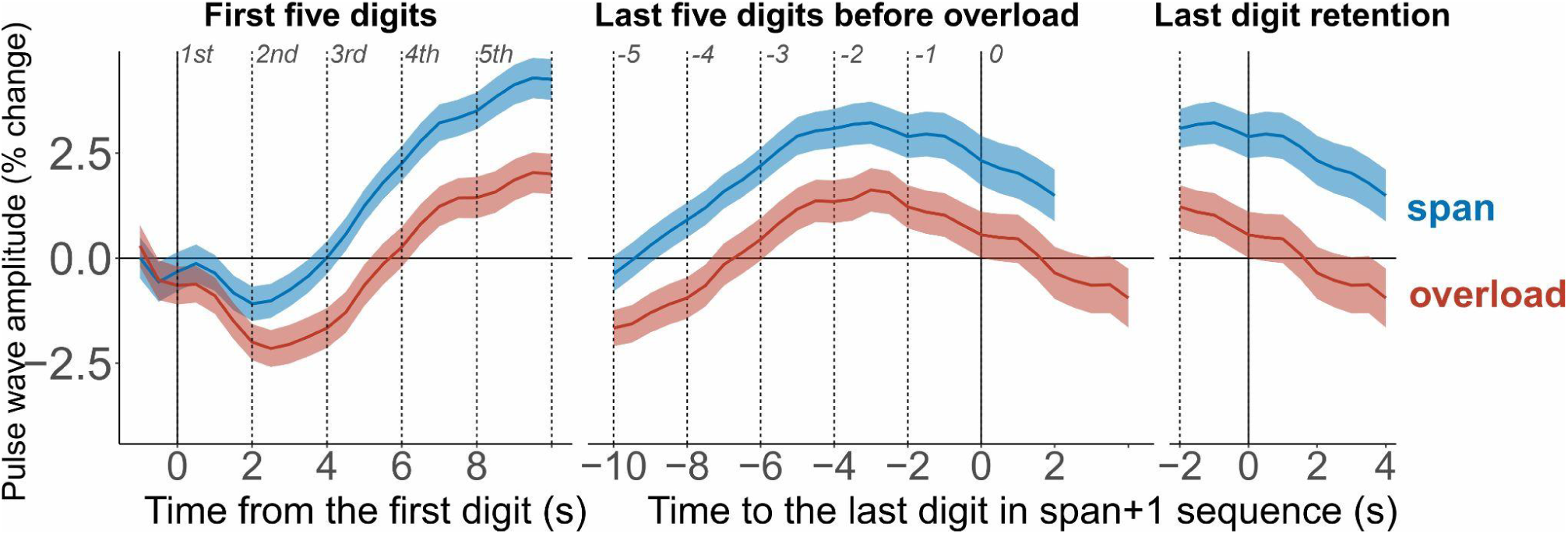
Pulse wave amplitude across sequence encoding and terminal encoding and retention in individual span and (span+1) overload conditions. Finger pulse wave amplitude (PWA) is expressed as percentage change from the 1-s presequence baseline. Blue denotes span and red denotes span+1. The left panel is aligned to the first digit. The middle panel is aligned to the additional final (span+1) overload digit (position 0); positions −5 to −1 correspond to the final five positions of the span sequence. The right panel realigns each condition to the onset of its own final digit to compare terminal encoding and retention. Vertical lines mark digit onsets. Baseline-normalized PWA was lower in overload (span+1) from the first digit onward and during terminal retention. Shaded bands show SEM.

#### First and last five digits

For the first five digits, PWA was generally lower in the overload condition than in the span condition (Condition: F(1, 116) = 18.16, p < .001, *η*_p_^2^ = .14; Serial position: F(1.66, 192.33) = 30.00, p < .001, *η*_p_^2^ = .21; Serial position × Condition: F(1.85, 214.53) = 7.60, p < .001, *η*_p_^2^ = .06). The difference between the span and overload conditions was already present in response to the first digit in the sequence and remained at a similar level across serial positions (1st: t(116) = 2.857, p = 0.005, d = 0.265; 2nd: t(116) = 3.522, p < 0.001, d = 0.327; 3rd: t(116) = 4.065, p < 0.001, d =0.377; 4th: t(116) = 3.968, p < 0.001, d = 0.368; 5th: t(116) = 4.003, p < 0.001, d = 0.372). Except for the first and third digit, all serial positions differed significantly from one another, with an initial decrease in baseline-normalized PWA from the first to the second digit, followed by a subsequent increase.

Similarly, PWA was lower in the overload condition in the last-five-digits comparison (Condition: F(1, 116) = 14.71, p < .001, *η*_p_^2^ = .113). PWA generally increased up to the second-to-last digit and then began to reverse, becoming lower toward the retention period (Serial position: F(1.52, 176.34) = 14.53, p < .001, *η*_p_^2^ =.11). However, this general pattern of PWA change did not differ between the overload and span conditions (Serial position × Condition: F(1.98, 230.02) = 0.23, p = .789, *η*_p_^2^ <.01). PWA increased in both the overload and span conditions up to the second-to-last digit, and the pattern reversed at the final digit. However, the difference between PWA at the second-to-last and final digit loads did not reach significance (p = 0.112). Similarly, the difference between the third- and second-to-last digit loads was not significant (p = 0.052), suggesting that PWA reached a plateau, whereas earlier pairwise comparisons showed a significant upward pattern.

### Tonic levels of raw pupil diameter and pulse wave amplitude

Raw PWA did not differ between span and span+1 during the 2-s pre-sequence period (t(118) = 0.86, p = .39, dz = 0.08) and did not differ during terminal encoding and retention either (t(118) = −1.41, p = .16, dz = 0.13; N = 119; Figure 6). Raw pupil diameter, in contrast, was larger in span+1 than in span in both intervals (pre-sequence period: t(108) = 5.25, p < .001, dz = 0.50; terminal encoding and retention: t(108) = 4.75, p < .001, dz = 0.45; N = 109). The tonic pupil difference was thus present before the sequence began and was removed by baseline correction, whereas the condition effects in the baseline-normalized PWA analyses did not reflect a difference in PWA level.

**Figure 6.**
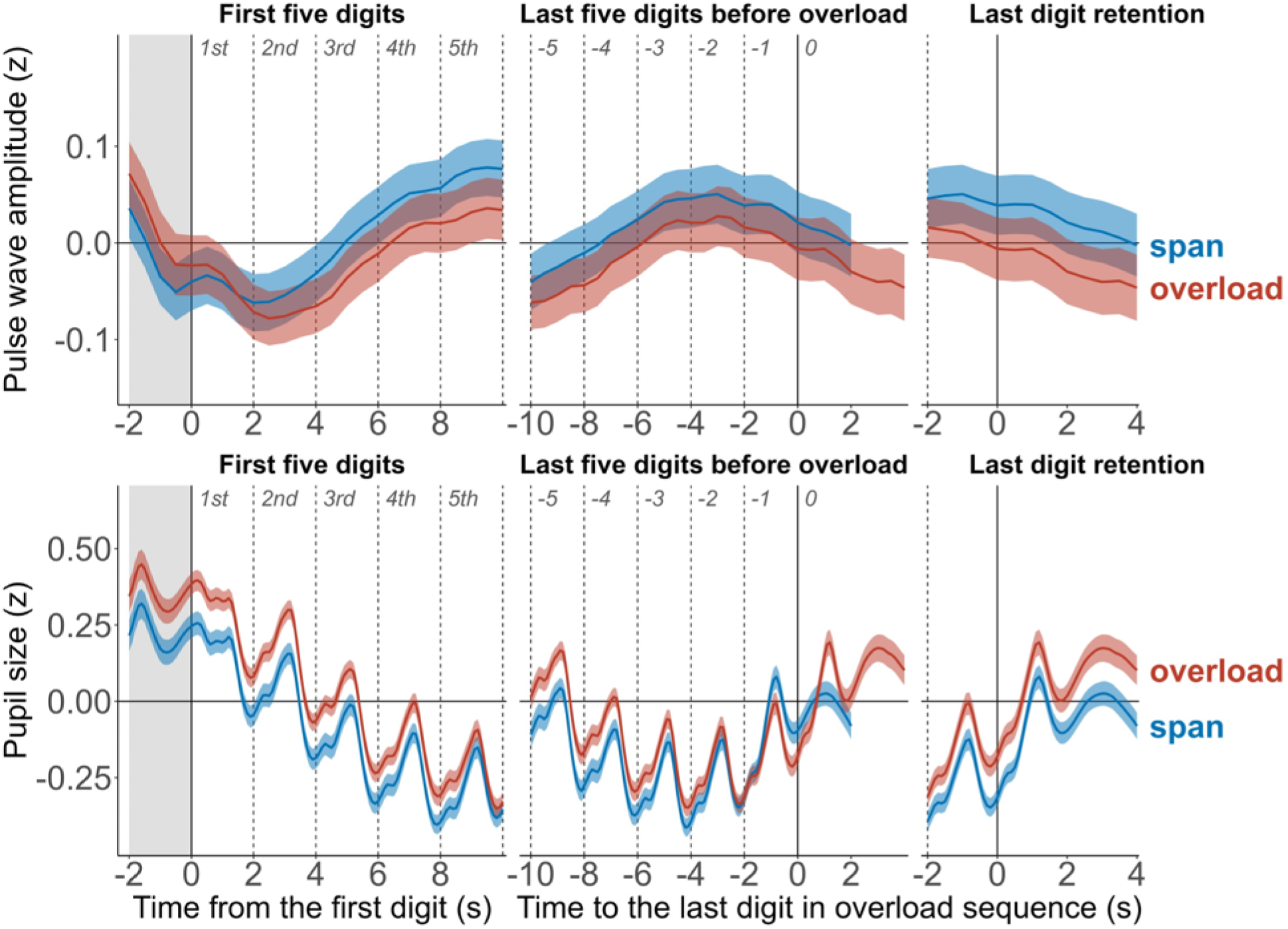
Non-baseline-corrected, within-participant standardized pulse wave amplitude (top) and pupil diameter (bottom) in individual span and span+1 overload conditions, z-scored within participant across all span and span+1 samples. As in Figures 4a and 5, the left panels are aligned to the first digit, the middle panels are aligned to the additional final span+1 digit at position 0, with −1 denoting the final span digit, and the right panels realign each condition to its own final digit. The grey band marks the 2-s pre-sequence period (−2 to 0 s), and dashed lines mark digit onsets. Blue denotes span and red denotes overload (span+1). Pupil diameter is larger in span+1 both before the sequence and during terminal encoding and retention, whereas PWA does not differ in either interval. Shaded bands show SEM.

### Electroencephalography

#### Span vs overload: retention interval

Theta increased more strongly during the last 4 s before the memory recall probe – during presentation of the last digit in the sequence and retention period – in the overload compared to span condition (F(1, 121) = 4.81, p = .030, *η*_p_^2^ = .04; see Figure 7). The same analysis did not reveal any differences in the alpha (F(1, 121) = 0.08, p = .780, *η*_p_^2^ < .01) or beta (F(1, 121) = 0.25, p = .621, *η*_p_^2^ < .01) frequency bands.

**Figure 7.**
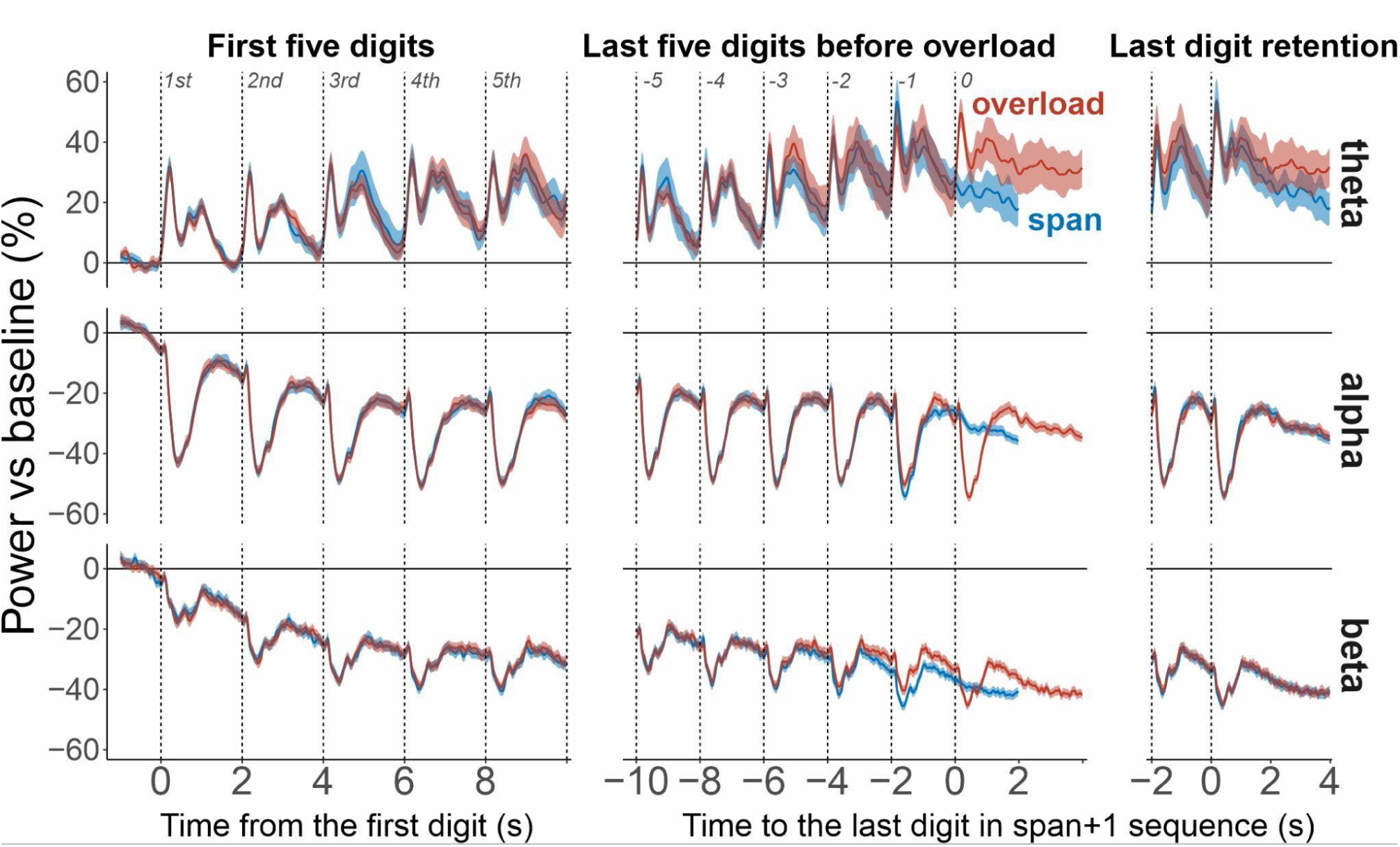
EEG power dynamics during encoding and retention of sequentially presented digits in individual span and overload (span+1) conditions. Grand average frontal midline theta (4-8 Hz; AFz, Fz, F1, and F2), posterior alpha (9-14 Hz; P1, P2, P3, P4, P5, P6, PO3, and PO4), and left centroparietal beta power (16-22 Hz; C3, C1, CP3, CP5, C5, and CP1) are expressed as percentage change from the 1-s presequence baseline. Blue denotes span and red denotes overload (span+1). The left panels are aligned to the first digit; the middle panels are aligned to the additional final span+1 digit at position 0, with −1 denoting the final span digit; and the right panels realign each condition to its own final digit. Vertical lines mark digit onsets. Theta increased with serial position and was greater during terminal encoding and retention in span+1. Alpha and beta decreased with serial position and showed condition differences only at late positions; neither differed between conditions in the terminal digit comparison. Shaded bands show SEM.

#### First and last five digits

For both the first and last 5 digits in the sequence, the effect of serial position was significant in all frequency bands (Table 2). Alpha and beta power decreased progressively with each added digit, whereas theta power increased.

**Table 2.**
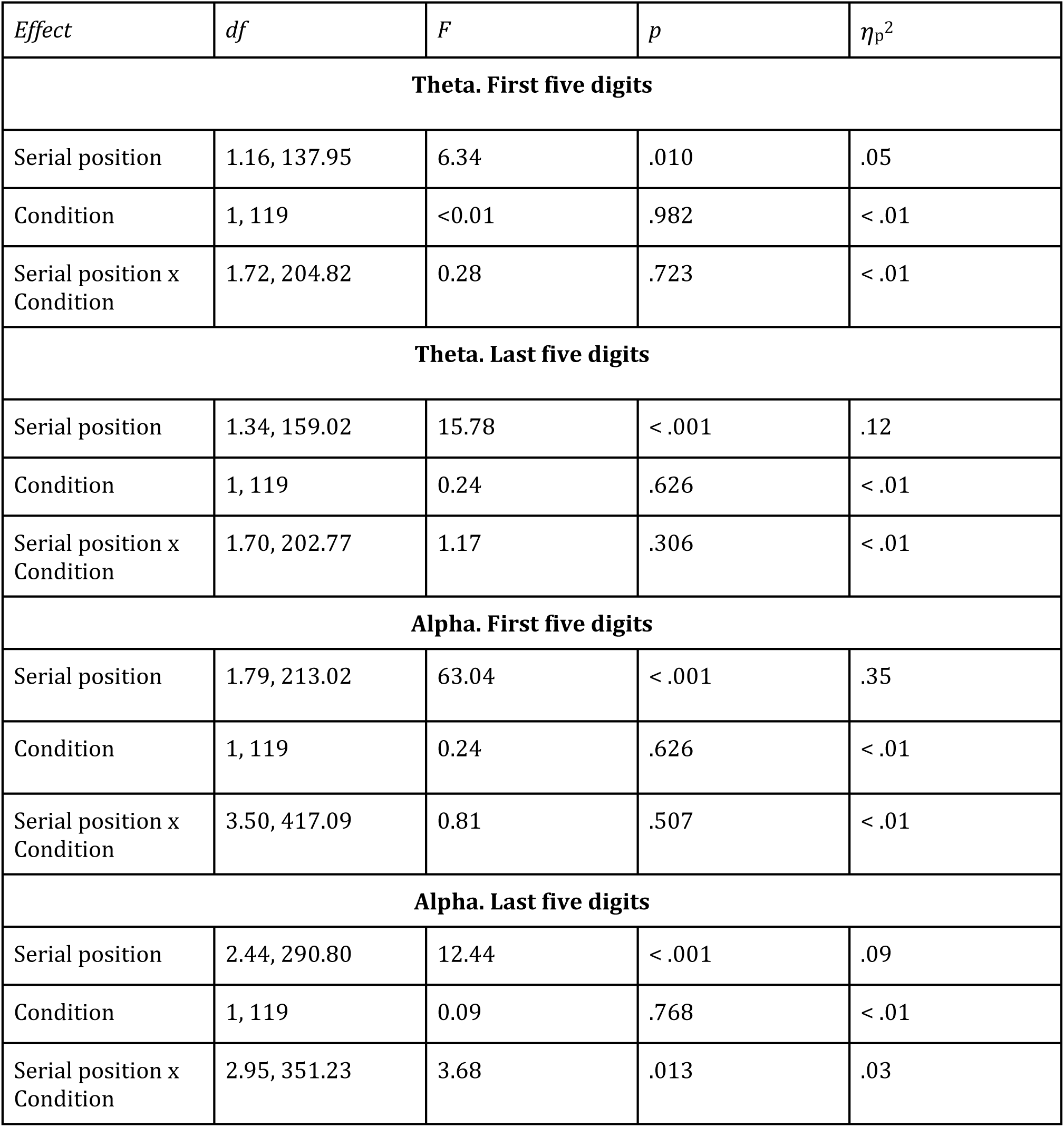
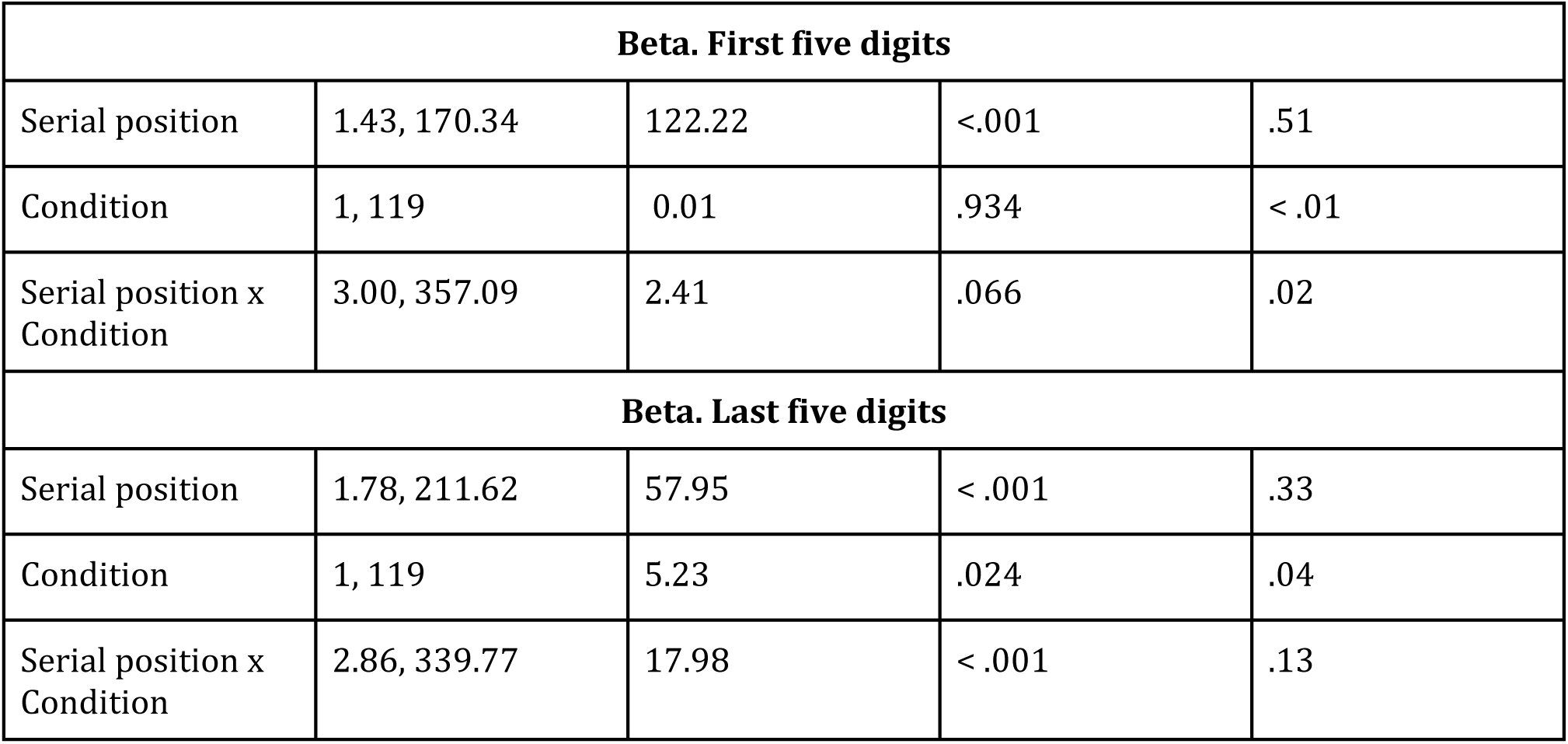
EEG ANOVA statistics for the first and last digits in the sequences.

| *Effect* | *df* | *F* | *p* | *η*<sub>p</sub><sup>2</sup> |
|---|---|---|---|---|
| **Theta. First five digits** |  |  |  |  |
| Serial position | 1.16, 137.95 | 6.34 | .010 | .05 |
| Condition | 1, 119 | <0.01 | .982 | < .01 |
| Serial position x Condition | 1.72, 204.82 | 0.28 | .723 | < .01 |
| **Theta. Last five digits** |  |  |  |  |
| Serial position | 1.34, 159.02 | 15.78 | < .001 | .12 |
| Condition | 1, 119 | 0.24 | .626 | < .01 |
| Serial position x Condition | 1.70, 202.77 | 1.17 | .306 | < .01 |
| **Alpha. First five digits** |  |  |  |  |
| Serial position | 1.79, 213.02 | 63.04 | < .001 | .35 |
| Condition | 1, 119 | 0.24 | .626 | < .01 |
| Serial position x Condition | 3.50, 417.09 | 0.81 | .507 | < .01 |
| **Alpha. Last five digits** |  |  |  |  |
| Serial position | 2.44, 290.80 | 12.44 | < .001 | .09 |
| Condition | 1, 119 | 0.09 | .768 | < .01 |
| Serial position x Condition | 2.95, 351.23 | 3.68 | .013 | .03 |
| **Beta. First five digits** |  |  |  |  |
| Serial position | 1.43, 170.34 | 122.22 | <.001 | .51 |
| Condition | 1, 119 | 0.01 | .934 | < .01 |
| Serial position x Condition | 3.00, 357.09 | 2.41 | .066 | .02 |
| **Beta. Last five digits** |  |  |  |  |
| Serial position | 1.78, 211.62 | 57.95 | < .001 | .33 |
| Condition | 1, 119 | 5.23 | .024 | .04 |
| Serial position x Condition | 2.86, 339.77 | 17.98 | < .001 | .13 |

For the first five digits in the sequence, no frequency band differentiated between the span and overload conditions, with no significant main effects of Condition and no significant interactions (see Table 2).

For the last five digits, there were no significant effects of Condition or Condition × Serial position interaction in the theta frequency band.

For alpha, although no significant main effect of Condition was present, the Condition × Serial position interaction was significant. This interaction indicated stronger alpha suppression in the span condition for the last analyzed digit in the sequence, i.e., the digit preceding the final digit in the overload condition and the actual final digit in the span condition (t(119) = 2.037, p = 0.044).

A similar interaction effect was observed in the beta frequency band, along with a main effect of Condition. The interaction suggested that, in the span condition, beta was more strongly suppressed in response to the final digit (t(119) = −5.029, p < .0001) and the second-to-last digit (t(119) = −2.794, p = .0061), but not to the other digits (third-to-last: t = 1.392, p = 0.1667; fourth-to-last: t = 0.101, p = 0.9199; fifth-to-last: t = 0.389, p = 0.6983). Thus, the two trajectories began to diverge earlier in the beta band than in the alpha band, but in a similar manner.

### Physiological differences associated with subsequent recall success

A separate analysis compared fully correct span+1 trials with trials containing at least one recall error (conveniently, it was an almost 50/50 split). For PWA and pupillometry, participants were required to contribute at least 10 usable trials; for EEG, they were required to contribute at least 15 clean trials. This analysis tested whether physiological activity during encoding and retention differentiated overload trials that were later recalled correctly from those containing an error.

PWA showed significant main effects of recall correctness in terminal encoding and retention interval (F(1, 114) = 9.11, p = .003, *η*_p_^2^ = .07; see Figure 8), the first-five-digit analysis (F(1, 113) = 5.36, p = .022, *η*_p_^2^ = .05), and the last-five-digit analysis (F(1, 112) = 5.85, p = .017, *η*_p_^2^ = .05) with lower PWA in correctly recalled trials.

**Figure 8.**
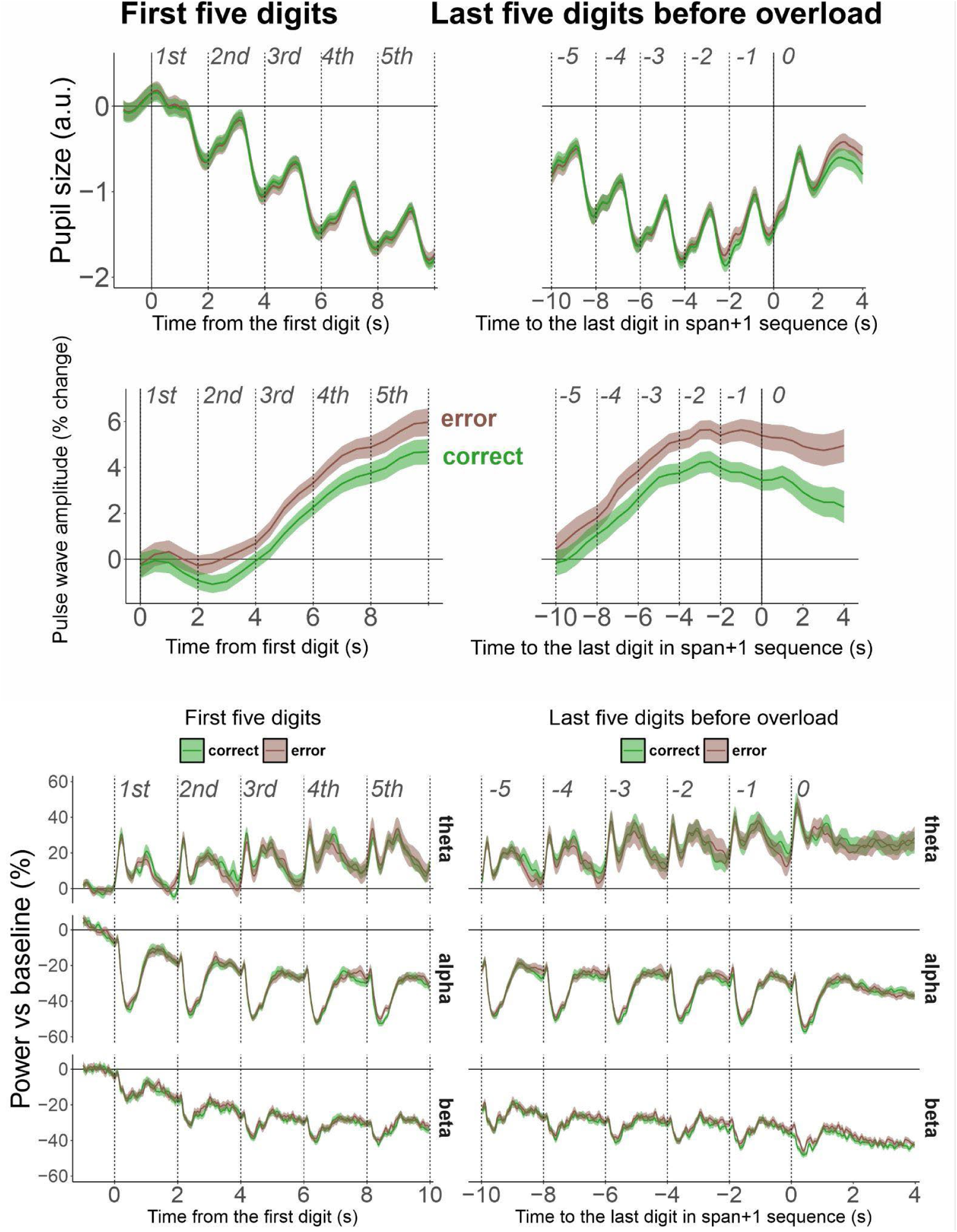
Physiological activity on correct and erroneous span+1 trials. Span+1 trials were classified as correct when the entire sequence was recalled in serial order and as erroneous when at least one digit was incorrectly recalled. Green denotes correct trials and brown denotes error trials. The left column is aligned to the first digit; the right column is aligned to the final span+1 digit at time 0. Vertical lines mark digit onsets. The upper row shows tonic pupil size in arbitrary units, the second row shows PWA as percentage change from baseline, and the lower rows show baseline-normalized frontal midline theta, posterior alpha, and left centroparietal beta power. PWA was more strongly suppressed on correct trials in the first-five, last-five, and terminal analyses. Pupil size did not differ by recall outcome. During terminal encoding and retention, beta suppression was stronger on correct trials, whereas theta and alpha did not differ. Shaded bands show SEM.

Pupil size did not differ by correctness during the terminal encoding and retention period (main effect of Correctness in retention period: F(1, 106) = 1.22, p = .272, *η*_p_^2^ = .01), during presentation of the first five digits (F(1, 105) = 0.08, p = .784, *η*_p_^2^ < .01), or last five digits (F(1, 104) = 0.08, p = .778, *η*_p_^2^ < .01). No interactions with serial position yielded significance in both measures.

During the terminal encoding and retention interval, beta power was more strongly suppressed in correctly recalled sequences (F(1, 94) = 8.41, p = .005, *η*_p_^2^ = .08). No corresponding effect of recall correctness was observed in alpha (F(1, 94) = 0.05, p = .828, *η*_p_^2^ < .01), or theta (F(1, 94) = 0.08, p = .777, *η*_p_^2^ < .01). Across the first- and last-five-digit EEG analyses, no main effects of correctness or Correctness × Serial Position interactions were significant in theta, alpha, or beta (Table S7).

## Discussion

We asked whether exceeding individually calibrated verbal working memory span by a single digit produces an identifiable change in neural or autonomic activity as the sequence unfolds. The span+1 manipulation increased subjective mental demand and substantially impaired recall: full-sequence accuracy decreased from 74.1% to 51.5%, and position-wise recall accuracy was lower throughout the sequence. Nevertheless, participants recalled considerable information under overload, including a slightly larger absolute number of digits in their correct positions (6.38 versus 6.12) despite lower proportional accuracy. Thus, the behavioral impairment did not resemble an abrupt loss of the ability to process additional items. The central physiological question was whether any recorded measure showed a corresponding transition when the added digit exceeded individual span.

Baseline-normalized pulse wave amplitude (PWA) was lower in the overload condition from the first digit onward and remained lower throughout the trial. A reduction in finger PWA is closely associated with sympathetically mediated peripheral vasoconstriction (Allen, 2007; Rahman et al., 2025; Zou et al., 2004). Similar relative attenuation of finger PWA has been reported during effortful cognitive tasks, including Sternberg working memory, mental arithmetic, and digit span task (Iani et al., 2004; Lyu et al., 2015; Pavlov et al., 2023). The difference between span and overload was already present at the first digit, well before an additional digit had been presented. Because participants knew the sequence length within each block, this early difference could reflect a tonic, block-level autonomic state. The raw-level analysis argues against this: raw PWA did not differ between span and span+1 during the 2-s pre-sequence period or during terminal encoding and retention. The condition differences in baseline-normalized PWA therefore do not reflect a tonic shift in peripheral vascular tone but a within-trial change of PWA relative to its pre-sequence level. In our previous fixed load study, PWA decreased with load but reversed at the highest sequence lengths (Pavlov et al., 2023). No comparable reversal was observed at individually calibrated span+1, and span+2 trials (see Supplementary materials) showed even lower baseline-normalized PWA. This pattern argues against the idea that the effort reflected in PWA is released as soon as span is exceeded. Across this individually calibrated overload range, PWA therefore tracked a sustained autonomic response to high task demands within each trial.

The supplementary span-1 comparison (see Supplementary materials) adds an important boundary related observation: PWA responses at span-1 and span were broadly similar, whereas the PWA was lower in span+1. This pattern is compatible with a change in autonomic responding around our operational overload boundary. Within span+1, PWA was also lower on subsequently correct than incorrect trials. Thus, PWA was associated both with the condition in which complete recall became unreliable and, within that condition, with whether recall succeeded. Because the group-level condition separation emerged before the additional digit, this subsequent-memory association is especially important: PWA appears to capture an aspect of the demanding task state that is relevant to performance, without identifying an error-causing event at a particular serial position.

Pupil size decreased during most of the trial and reversed only near the final digit in the sequence. This differs from the typical pattern of increasing pupil dilation with each additional memory item, followed by a plateau or decline when demands exceed capacity (Cabestrero et al., 2009; Granholm et al., 1996; Johnson et al., 2014; Kosachenko et al., 2023). Unlike those studies using auditory presentation, we presented white visual digits against a gray background. Stimulus luminance can substantially reduce or even reverse task-evoked pupil dilation (Kay et al., 2022; Klingner et al., 2011; Pan et al., 2022, 2024; Peysakhovich et al., 2017). Thus, the overall pupil size dynamics were likely influenced by both cognitive load and stimulus luminance. This downward trend was eliminated in the analysis of phasic pupil responses, in which pupil size was time-locked to individual digit presentation and baseline-corrected before each digit. In that analysis, each newly added digit produced a reliable increase in pupil size relative to the preceding digit, which is consistent with the expected working memory load effect.

The condition difference in pupil size was temporally specific and similar in tonic and phasic pupil response analyses. Pupil size did not differ reliably between span and overload during the first five digits or during the final retention interval when each condition was aligned to its own final digit. The difference appeared at late positions matched to the end of the span sequence: tonic pupil size was smaller in overload at the last three matched digits, and phasic responses were smaller at the final two matched positions. This pattern appears to reflect a temporal shift in pupil size dynamics: pupil size began to increase earlier in the span condition and later in the overload condition. One possibility is that participants modulated their effort during late encoding in anticipation of the extra demand, producing a transient attenuation of pupil responses. The convergence of phasic responses at the respective final digits suggests that this modulation was temporary. This interpretation is consistent with the broader view that task-evoked pupil responses index effort allocation and control demands, but are not a pure measure of objective task difficulty or memory load (Beatty, 1982; Mathôt, 2018; Van Der Wel & Van Steenbergen, 2018).

The baseline-corrected analyses, however, conceal a tonic effect. Raw pupil diameter was larger in span+1 than in span blocks both during the 2-s pre-sequence period and during terminal encoding and retention, whereas the baseline-corrected retention comparison showed no difference. Because the sequence length was known before each block, this tonic dilation most plausibly reflects anticipatory arousal in the more demanding blocks, consistent with the role of tonic pupil size as an index of locus coeruleus-mediated arousal (Aston-Jones & Cohen, 2005; Mathôt, 2018). The temporal shift in baseline-corrected responses described above therefore occurred on top of a tonically enlarged pupil.

The different temporal dynamics of PWA and pupil responses suggest that autonomic responses to working memory overload involve distinct processes. PWA primarily reflects sympathetically driven peripheral vascular tone. Pupil diameter reflects a more complex combination of the pupillary light reflex, parasympathetic inhibition of the iris sphincter, and sympathetic input to the iris dilator (Aston-Jones & Cohen, 2005; Steinhauer et al., 2004; Szabadi, 2012). The analyses of non-baseline-corrected pupil size and PWA sharpen this dissociation: pupil diameter, but not PWA, was tonically larger in overload blocks, both before the sequence began and during retention. One possibility is that tonic pupil dilation reflects a central arousal state set by the expected difficulty of the block, whereas the peripheral vascular response unfolds within each trial in proportion to the demands of the sequence. Thus, mild overload involved a tonic increase in pupil-indexed arousal together with a within-trial, but not tonic, peripheral vascular response.

Posterior alpha was sensitive to cognitive load, but it did not provide a simple marker of overload. We observed a load-dependent alpha decrease, without a clear load-dependent phasic alpha enhancement after the switch from encoding (0.5 s) to the retention period following each digit presentation (1.5 s), as would have been predicted by some accounts of inhibition of irrelevant sensory processing (Jensen & Mazaheri, 2010). In the present task, each visually presented digit had to be encoded, integrated into serial order, and maintained for later typing. Progressive alpha suppression is therefore consistent with sustained visual and attentional engagement as the sequence grows (Jensen & Mazaheri, 2010; Klimesch, 2012; Klimesch et al., 2007).

Alpha was sensitive to the sequence stage. At the position corresponding to the final digit of the span sequence, alpha was more strongly suppressed in span than in overload, where one additional digit remained. This pattern suggests that alpha activity was also sensitive to the changing demands of the final stages of encoding. The actual final digit may receive greater attentional processing as the complete sequence becomes available for maintenance and subsequent recall. The stronger suppression in span is consistent with such increased processing of the terminal item, whereas overload required continued incorporation of another digit. Stronger alpha suppression at this point in the span condition is therefore consistent with an end-of-encoding transition, similar to the dynamics observed for pupil size, possibly involving retrieval preparation, reorganization of the maintained sequence, or increased attentional engagement with the final item.

A related but more pronounced late effect was observed in beta activity. Sensorimotor beta suppression is commonly linked to motor preparation and action planning (Boettcher et al., 2021; Kosachenko et al., 2026; Schneider et al., 2017; van Wijk et al., 2012). Because participants recalled the sequence by typing the digits in order, accumulating memory representations could also support the gradual formation of an ordered response plan. This interpretation is consistent with studies showing that working memory representations are often coupled to anticipated actions (Olivers & Roelfsema, 2020; Rösner et al., 2022; Van Ede, 2020), and with studies linking beta modulation to motor sequence learning and explicit sequence knowledge (Dyck & Klaes, 2024; Meissner et al., 2018; Pollok et al., 2014). Beta suppression was stronger at the last two span-matched positions in span than in overload, consistent with earlier preparation of the forthcoming typed response when the end of the sequence was closer. The span-1 comparison (see Supplementary materials) also showed an orderly late divergence across sequence lengths, supporting sensitivity to the stage of encoding. Of particular interest, during terminal encoding and retention beta suppression was stronger on correctly recalled than incorrectly recalled span+1 trials. This association links late beta activity to the successful transformation of maintained information into a response, making the action-oriented interpretation a promising account of the observed dynamics. An alternative, not mutually exclusive, account is subvocal rehearsal: the left centro-parietal cluster overlaps sensorimotor regions in which mu/beta suppression accompanies covert as well as overt articulation (Bowers et al., 2019; Jenson et al., 2014; Saltuklaroglu et al., 2018), and rehearsal would also be expected to intensify as the list grows and to differ between correct and erroneous trials. The present data cannot distinguish rehearsal from response preparation, since both engage left sensorimotor cortex and both scale with sequence length.

Frontal midline theta followed a different pattern. Theta power increased with serial position and was greater during terminal encoding and retention in span+1 than in span. This result is consistent with extensive evidence linking frontal theta to cognitive control demands and working memory (Cavanagh & Frank, 2014; Pavlov & Kotchoubey, 2022; Roux & Uhlhaas, 2014). As more digits were presented, successful performance likely required increasing control over the maintenance of serial order and interference from previously encoded items. The continued theta increase suggests that these control-related processes remained active as participants approached and even exceeded individual span. Interestingly, theta and phasic pupil responses initially showed similar load-dependent increases, but their dynamics diverged during late encoding. Whereas phasic pupil responses were attenuated in overload at the last matched serial positions and converged at the respective final digits, theta showed no corresponding attenuation and was greater during terminal encoding and retention in overload. This pattern is consistent with continued, and possibly compensatory, recruitment of cognitive control in an attempt to stabilize performance under excessive memory demands. The span-1 analysis (see Supplementary materials) supports this interpretation: theta was lower below span, but span and span+1 were more similar, suggesting that control demands were already strongly elevated at the capacity boundary and remained high under overload. In this respect, theta differed from pupil responses, which showed a transient attenuation during late encoding. The two measures therefore appear to capture different aspects of effortful performance: frontal theta more directly tracked increasing control demands, whereas pupil responses were more sensitive to changes in effort allocation as overload was anticipated.

## Conclusion

By calibrating memory load to each participant’s span in a large sample with simultaneous EEG, pupillometry and photoplethysmography, we characterized what happens when working memory capacity is exceeded by a single item. Overload reduced full-sequence recall from 74% to 52% and added little to the number of digits recalled in their correct positions, yet it produced no abrupt collapse or discontinuity in neural or autonomic activity. Meanwhile, overload produced three dissociable physiological responses. First, an anticipatory response: raw pupil diameter was larger in overload blocks before the first digit appeared and remained larger during retention. Second, a trial-level effort response: pulse wave amplitude differed between span and overload conditions – within each trial it was attenuated from the first digit onward, more so at span+2 and on overload trials that were later recalled correctly. Third, end-of-list processing: alpha and beta power diverged mainly toward the end of encoding, and terminal beta suppression also predicted recall success. Frontal midline theta, meanwhile, kept rising with the number of items held.

At the limit of capacity, then, people neither disengage nor stop encoding the final items, and cognitive control related activity does not collapse. Effort is mobilized in anticipation of the harder block and maintained, even increased, within each trial as the chance of success falls. Motivational intensity theory predicts exactly this for as long as success remains possible (Brehm & Self, 1989; Gendolla & Richter, 2010). Reversals of physiological responses at high loads in fixed-load designs are not associated with a mere exceedance of capacity limits but only disengagement due to facing an impossible task leading to a certain failure, which makes individual calibration essential for studying overload. Methodologically, our results caution that per-condition baseline correction can hide anticipatory effects in blocked designs. Practically, PWA, measurable with a simple finger sensor, tracked both individualized overload and recall success, making it a promising candidate for personalized cognitive load monitoring.

Future work should identify which process fails on the roughly half of overload trials that end in error. Two analyses could show whether overload degrades item representations or their binding to serial positions: separating item from order errors in the recall strings, and decoding item and position information from multivariate EEG. Both are possible in the openly available present dataset. Sequences of unannounced length would separate anticipatory from load driven responses. Extending individualized exceedance beyond span+2, with ratings of expected success, would test where effort is withdrawn. Finally, auditory or isoluminant presentation would allow phasic pupil responses to be measured without the luminance confound of visual digits.

## Author contributions statement

Alexander I. Kotyusov: Investigation, Methodology, Formal analysis, Data curation, Writing - original draft, Writing - review & editing.

Dauren Kasanov: Investigation, Methodology, Data curation, Writing - review & editing. Alexandra I. Zaitseva: Investigation, Methodology, Data curation, Writing - review & editing. Boris Kotchoubey: Writing - review & editing.

Yuri G. Pavlov: Conceptualization, Funding acquisition, Data curation, Formal analysis, Project administration, Supervision, Visualization, Methodology, Writing - original draft, Writing - review & editing.

## Data availability statement

The data are publicly available on OpenNeuro (https://openneuro.org/datasets/ds008302).

## Potential Conflicts of Interest

Nothing to report.

## Acknowledgements

Authors AIK, DK, AIZ were supported by RSF grant 25-78-10176

## Supplement

### Above overload: span+2

To examine physiology under a more pronounced overload, we conducted supplementary analyses of the span+2 condition. The span+2 analyses also aimed to address whether the span+1 condition represented a sufficiently strong capacity exceedance via comparing overall trends in three conditions directly: span, span+1, span+2.

Because not all participants reached this level, the samples were necessarily smaller than in the primary analyses. For these supplementary analyses, we required at least 10 usable trials for the PWA and pupillometry analyses and at least 15 trials for the EEG analyses.

### Behavior

A total of 88 participants contributed at least 10 span+2 trials. The mean proportion of fully recalled sequences fell from span to span+1 to span+2 (F(1.74, 151.70) = 298.46, p < .001, *η*_p_^2^ = .77; 73.7%, 54.7% and 32.7%). Every pair of conditions differed (all p < .001; dz = 1.38–2.26). Partial accuracy showed the same pattern (F(1.54, 133.87) = 162.53, p < .001, *η*_p_^2^ = .65; 90.2%, 83.4% and 73.4%; all pairs p < .001). Mental demand rose step by step (F(1.39, 101.71) = 138.15, p < .001, *η*_p_^2^ = .65; 10.7, 12.6 and 14.3; all pairs p < .001). The partial score (number of digits recalled) was higher in span+1 and span+2 than in span (F(1.43, 124.83) = 16.82, p < .001, *η*_p_^2^ = .16; both p < .001). Span+1 and span+2 did not differ on this measure (6.52 vs 6.51, p = .907), so the extra digit added no more recalled digits.

**Table S1.** Behavioral accuracy and subjective cognitive load in span+2 condition.

| Condition | Mental demand, M (SD) | Partial-recall accuracy, %, M (SD) | Digits correct, n, M (SD) | Median sequence length | Correct sequences, %, M (SD) |
|---|---|---|---|---|---|
| Span+2 | 14.3 (3.26) | 73.4 (9.84) | 6.51 (1.56) | 9 | 32.7 (15.3) |

### Pupil

Eighty-two participants met the inclusion criterion for the span+2 pupillometry analyses.

Tonic pupil size during presentation of the final digit and the subsequent retention interval did not differ across the three sequence-length conditions (F(1.47, 118.92) = 1.11, p = .318, *η*_p_^2^ = .01; see Figure S1A).

Across the first five digits, tonic pupil size changed with serial position, showing the overall downward trajectory observed in the primary analysis (Serial position: F(1.39, 109.52) = 72.79, p < .001, *η*_p_^2^ = .48). On average the conditions did not differ (main effect of Condition: F(1.69, 133.61) = 0.31, p = .700, *η*_p_^2^ < .01), and there was no significant Serial Position × Condition interaction (F(3.87, 305.48) = 0.85, p = .491, *η*_p_^2^ = .01).

In the analysis of the last five span-matched digits, tonic pupil size showed significant effects of Condition (F(1.57, 123.93) = 3.48, p = .045, *η*_p_^2^ = .04), Serial position (F(1.19, 93.98) = 23.94, p < .001, *η*_p_^2^ = .23), and their interaction (F(4.08, 322.16) = 14.03, p < .001, *η*_p_^2^ = .15). At the final matched position, corresponding to the final digit of the span sequence, all three pairwise contrasts were significant (span > span+1: p<0.001; span > span+2: p<0.001; span+1 > span+2: p=0.0177). At the second-to-last matched position, span differed from span+1 (p = .0164) and span+2 (p = .010), whereas span+1 and span+2 did not differ (p = .387).

For the first five digits, phasic pupil responses increased with serial position (Serial position: F(2.90, 228.97) = 14.63, p < .001, *η*_p_^2^ = .16). This increase did not differ by Condition (main effect of Condition: F(1.68, 132.53) = 0.13, p = .840, *η*_p_^2^ < .01; Serial Position × Condition: F(6.01, 474.72) = 1.99, p = .066, *η*_p_^2^ = .02).

In the analysis of the last five span-matched digits, phasic pupil responses again increased with Serial position (F(3.55, 280.21) = 25.25, p < .001, *η*_p_^2^ = .24). The main effect of condition was significant (F(1.65, 130.34) = 6.01, p = .006, *η*_p_^2^ = .07) with responses being larger in span than in span+1 (p < .001) and span+2 (p = .008), whereas span+1 and span+2 did not differ (p = .619). The Serial Position × Condition interaction was also significant (F(5.50, 434.60) = 6.32, p < .001, *η*_p_^2^ = .07).

Post hoc comparisons localized this interaction (see Table S2). At the fifth-to-last position, phasic pupil responses were larger in span+2 than in span and span+1, whereas span and span+1 did not differ. No pairwise differences were observed at the fourth-to-last or third-to-last positions. At the second-to-last position, span responses exceeded span+1 responses, whereas the other contrasts were not significant. At the final matched position, all three contrasts were significant. Despite these late differences, the final digit response itself converged across conditions (F(1.72, 139.65) = 0.29, p = .715, *η*_p_^2^ < .01; Figure S1B).

**Table S2.** Pairwise post hoc comparisons of phasic pupil responses by span sequence length aligned serial position.

| Serial position | Contrast | df | t | p |
|---|---|---|---|---|
| -5 | Span - Span + 1 | 79 | 0.36 | .7200 |
|  | Span - Span + 2 | 79 | -2.49 | .0149 |
|  | Span + 1 - Span + 2 | 79 | -2.69 | .0087 |
| -4 | Span - Span + 1 | 79 | 0.68 | .4969 |
|  | Span - Span + 2 | 79 | 1.05 | .2975 |
|  | Span + 1 - Span + 2 | 79 | 0.73 | .4701 |
| -3 | Span - Span + 1 | 79 | -1.26 | .2110 |
|  | Span - Span + 2 | 79 | -0.42 | .6774 |
|  | Span + 1 - Span + 2 | 79 | 0.53 | .5973 |
| -2 | Span - Span + 1 | 79 | 2.37 | .0202 |
|  | Span - Span + 2 | 79 | 0.51 | .6105 |
|  | Span + 1 - Span + 2 | 79 | -1.08 | .2847 |
| -1 | Span - Span + 1 | 79 | 4.41 | < .001 |
|  | Span - Span + 2 | 79 | 4.77 | < .001 |
|  | Span + 1 - Span + 2 | 79 | 2.74 | .0077 |

*Note*. Serial position is coded relative to the final item of the span+1 sequence. Positions -5 to -1 correspond to the last five positions of the span sequence; position -1 is the final digit of the span sequence, and position 0 would be the final digit of the span+1 sequence (not included in this analysis). Estimates are calculated as the first condition minus the second condition in each contrast.

**Figure S1.**
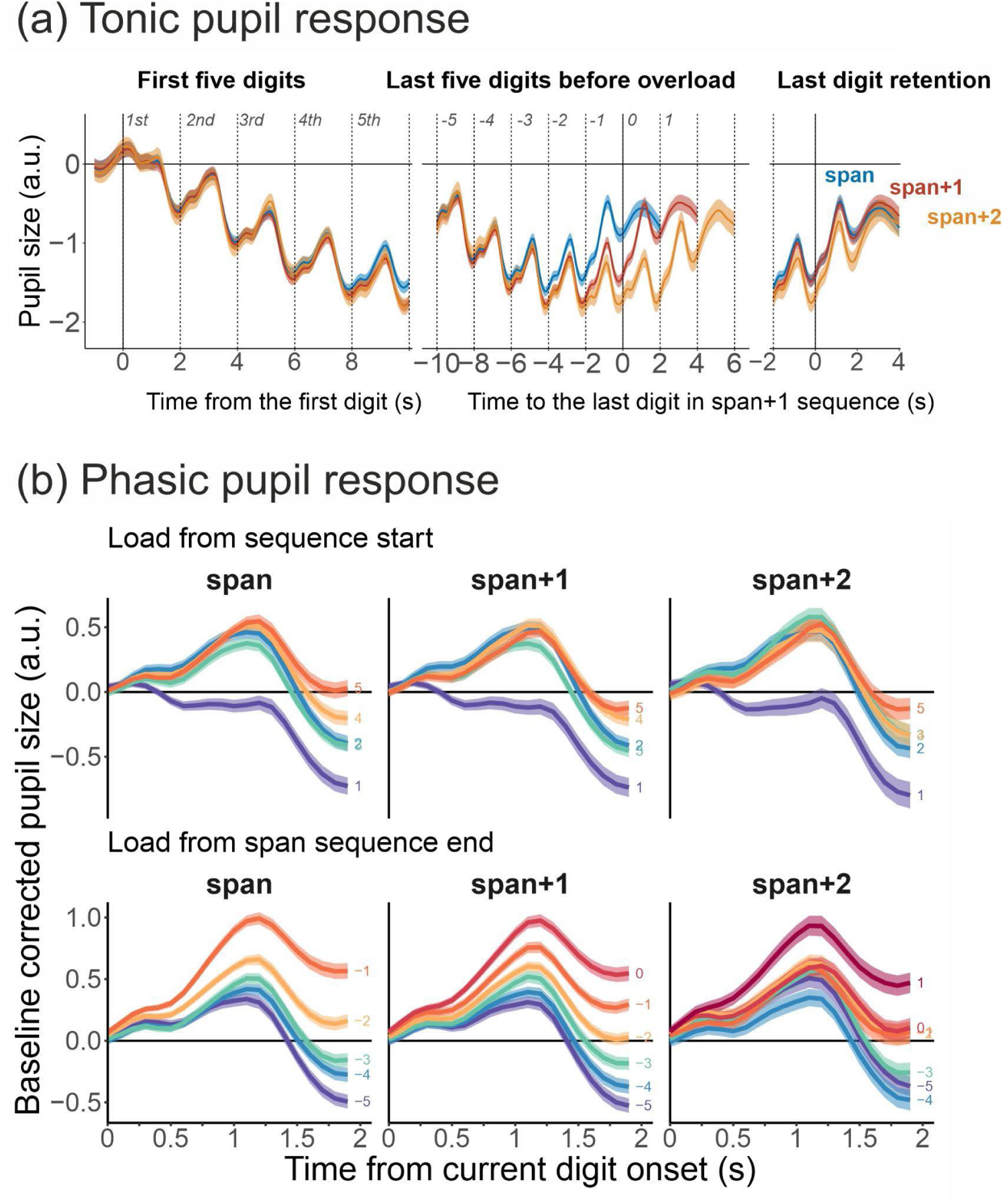
Tonic and phasic pupil responses at span, span+1, and span+2. Blue denotes span, red denotes span+1, and orange denotes span+2. The upper row shows tonic pupil size after subtraction of the 1-s presequence baseline, aligned to sequence onset, to the final span+1 digit, and to each condition’s own final digit. The lower rows show phasic responses baseline-corrected to the 200 ms immediately preceding each digit. Positions 1-5 are counted from sequence onset. Late positions are coded relative to the final span+1 digit: −1 is the final span digit, 0 is the final span+1 digit, and +1 is the final span+2 digit. Phasic responses increased across serial positions, differed among conditions at selected late positions, and converged at the respective final digit. Shaded bands show SEM.

### PWA

Eighty-eight participants met the inclusion criterion of at least 10 usable trials for the span+2 PWA analyses.

During the final retention interval, PWA was more strongly attenuated as sequence length increased (F(1.52, 132.41) = 21.49, p < .001, *η*_p_^2^ = .20; see Figure S2).

**Figure S2.**
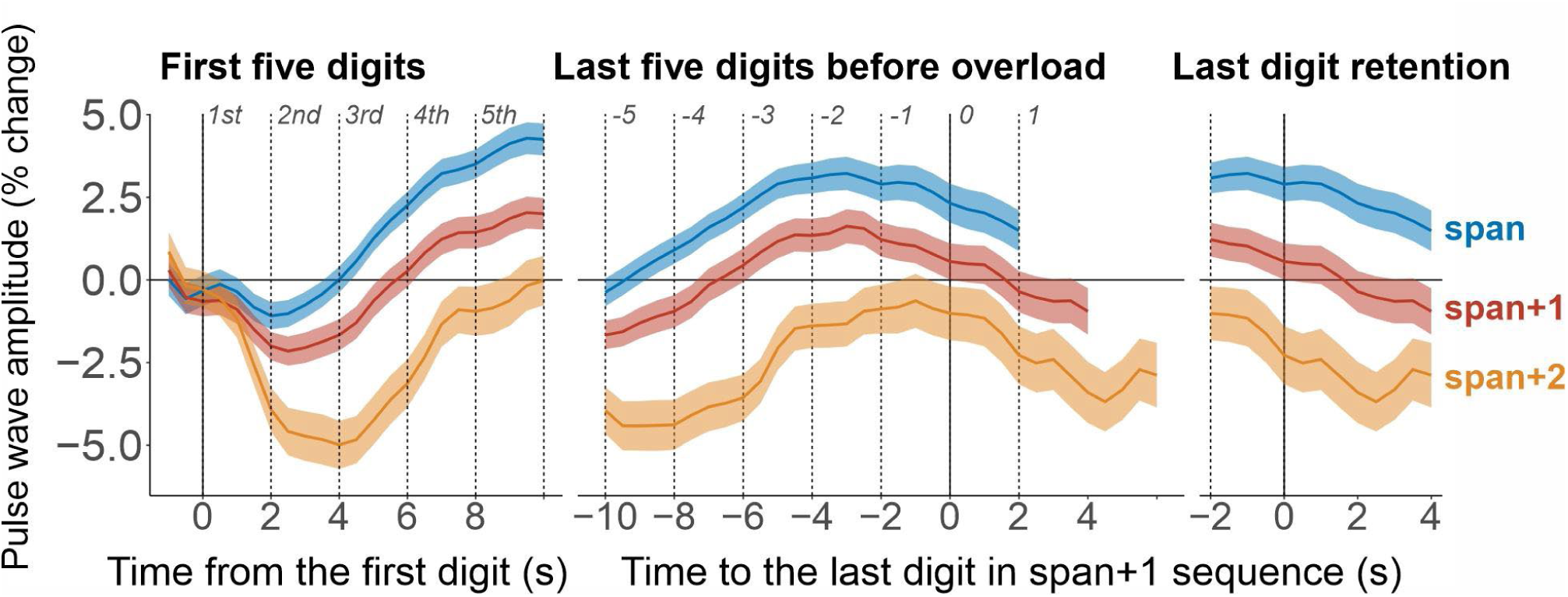
Pulse wave amplitude at span, span+1, and span+2. PWA is expressed as percentage change from the 1-s presequence baseline. Blue denotes span, red denotes span+1, and orange denotes span+2. The left panel is aligned to sequence onset. The middle panel is aligned to the final span+1 digit: −1 is the final span digit, 0 is the final span+1 digit, and +1 is the final span+2 digit. The right panel realigns each condition to its own final digit. Vertical lines mark digit onsets. PWA was progressively lower from span to span+1 to span+2 during both early and late encoding and during terminal encoding and retention. Shaded bands show SEM.

Across the first five digits, the PWA attenuation observed in span+1 was replicated. It was stronger in span+2 than in both span and span+1 (main effect of Condition: F(1.78, 150.92) = 40.87, p < .001, *η*_p_^2^ = .32) The effect of Serial position (F(1.77, 150.47) = 15.65, p < .001, *η*_p_^2^ = .16) and Serial position x Condition interaction (F(3.18, 270.36) = 12.22, p < .001, *η*_p_^2^ = .13) were significant. All pairwise comparisons between conditions were significant at every serial position. The interaction indicated that the condition effect was somewhat smaller at the first digit, but remained significant at each position.

The analysis of the last five digits showed the same rank order, with PWA highest in span, lower in span+1, and lowest in span+2 (F(1.74, 147.67) = 36.29, p < .001, *η*_p_^2^ = .30). PWA also varied with serial position (F(1.48, 125.46) = 10.89, p < .001, *η*_p_^2^ = .11), whereas the Serial Position × Condition interaction was not significant F(2.76, 235.00) = 2.39, p = .075, *η*_p_^2^ = .03).

### EEG

Forty-six participants remained in the EEG analyses after applying the criterion of at least 15 clean span+2 trials. For comparison, the primary span and span+1 EEG analyses included N = 122 participants.

During the terminal encoding and retention interval, theta did not show a further significant increase in span+2 compared to span+1 (p = 0.356), although the difference between span and span+2 was significant (p = 0.014) (main effect of Condition: F(2, 90) = 3.93, p = .023, *η*_p_^2^ = .08). The condition effect was significant for beta (F(2, 90) = 5.33, p = .007, *η*_p_^2^ = .11) due to weaker beta suppression in span+2 than in span (p = 0.007) and span+1 (p = 0.007). The corresponding condition effect in alpha did not reach significance (F(1.51, 67.80) = 2.83, p = .080, *η*_p_^2^ = .06).

**Table S3.** EEG ANOVA statistics for serial position and condition across span, span+1, and span+2 trials.

| *Effect* | *df* | *F* | *p* | *η*<sub>p</sub><sup>2</sup> |
|---|---|---|---|---|
| **Theta. First five digits** |  |  |  |  |
| Serial position | 2.32, 102.11 | 5.80 | .003 | .12 |
| Condition | 1.35, 59.34 | 1.19 | .296 | .03 |
| Serial position x Condition | 3.48, 153.07 | 1.49 | .214 | .03 |
| **Theta. Last five digits** |  |  |  |  |
| Serial position | 1.54, 67.97 | 8.04 | .002 | .15 |
| Condition | 1.38, 60.75 | 0.77 | .424 | .02 |
| Serial position x Condition | 3.63, 159.83 | 0.80 | .519 | .02 |
| **Alpha. First five digits** |  |  |  |  |
| Serial position | 1.51, 66.55 | 11.06 | < .001 | .20 |
| Condition | 1.77, 77.87 | 2.30 | .114 | .05 |
| Serial position x Condition | 5.23, 230.33 | 1.02 | .410 | .02 |
| **Alpha. Last five digits** |  |  |  |  |
| Serial position | 2.45, 107.61 | 0.78 | .485 | .02 |
| Condition | 2, 88 | 2.82 | .065 | .06 |
| Serial position x Condition | 4.68, 205.77 | 1.86 | .107 | .04 |
| **Beta. First five digits** |  |  |  |  |
| Serial position | 1.49, 65.74 | 26.83 | < .001 | .38 |
| Condition | 1.75, 77.00 | 1.67 | .198 | .04 |
| Serial position x Condition | 4.92, 216.36 | 0.84 | .521 | .02 |
| **Beta. Last five digits** |  |  |  |  |
| Serial position | 1.82, 80.08 | 4.83 | .013 | .10 |
| Condition | 2, 88 | 3.93 | .023 | .08 |
| Serial position x Condition | 3.20, 140.88 | 6.01 | < .001 | .12 |

**Figure S3.**
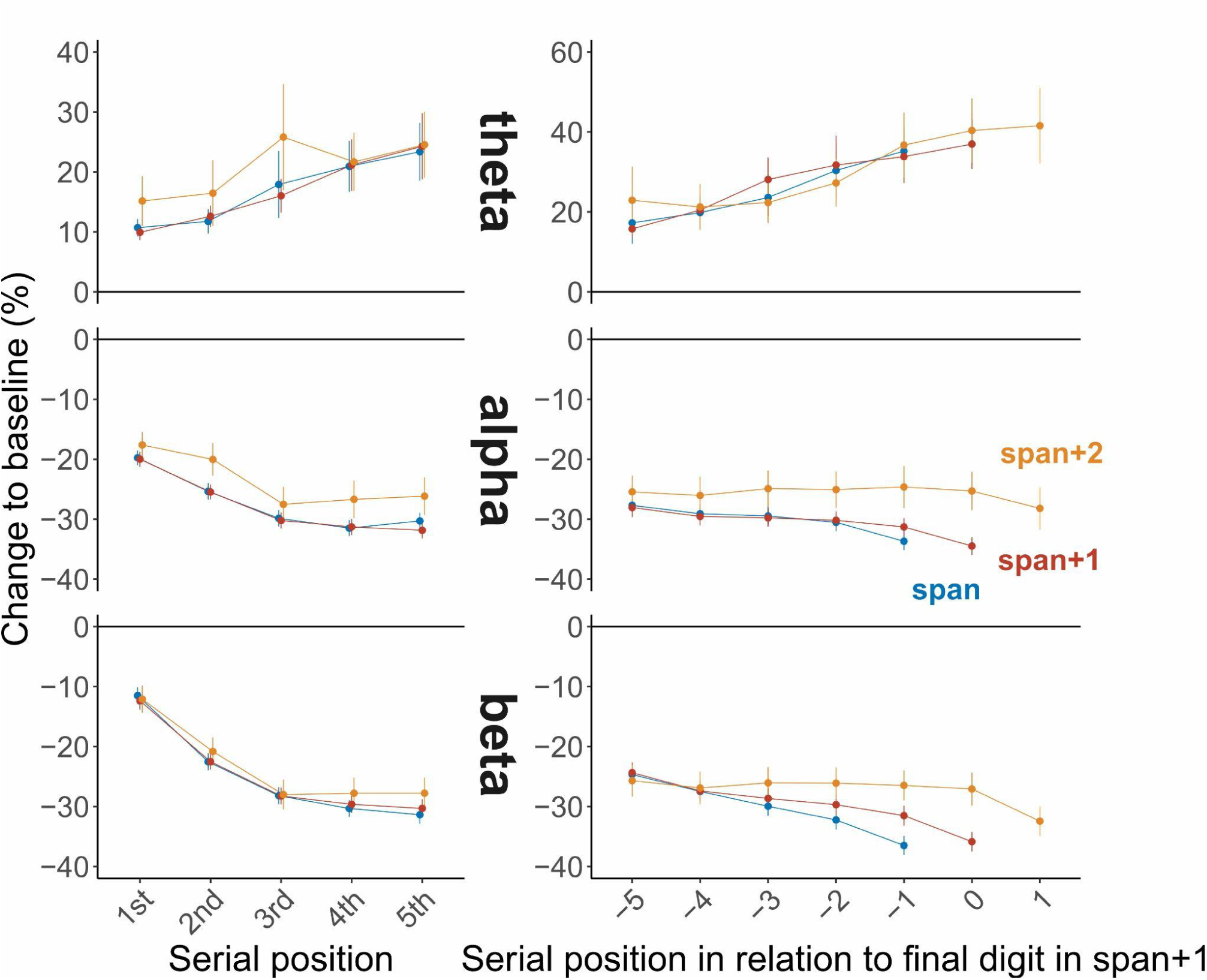
Theta, alpha, and beta power across span, span+1, and span+2. Group means are shown for frontal midline theta, posterior alpha, and left centroparietal beta power, expressed as percentage change from the presequence baseline. Blue denotes span, red denotes span+1, and orange denotes span+2. The left column shows positions 1-5 from sequence onset. In the right column, positions are coded relative to the final span+1 digit: −1 is the final span digit, 0 is the final span+1 digit, and +1 is the final span+2 digit. Theta increased and alpha and beta decreased across the first five positions, with no significant condition effects or interactions. During late encoding, beta showed significant effects of condition and a Condition × Serial Position interaction; corresponding condition effects were not significant for theta or alpha. Error bars show SEM.

Across the first five digits, the three condition analysis reproduced the principal serial position effects observed in the primary analysis: theta increased with serial position, whereas alpha and beta decreased (Table S3; Figure S3). No main effects of Condition or Serial Position × Condition interactions reached significance in any frequency band during this portion of the sequence.

While expectedly we observed a generally noisier pattern due to fewer trial averages and fewer remaining participants, in the last five digit analysis, theta again changed with serial position, but neither the main effect of Condition nor the interaction was significant. Alpha showed no significant Serial position effect, no Condition effect, and no interaction. Beta showed significant effects of serial position: the late beta trajectory differed across span, span+1, and span+2, with weaker beta suppression in span+2.

### Easier than at capacity: span-1

To test whether overload marks a quantitatively distinct physiological state, which would differ from a continuation of the linear physiological parameter change trajectory, we compared the span-1, span, and span+1 conditions. If span+1 reflects a distinct overload state, span-1 and span should be relatively similar because both remain within capacity, whereas span+1 should diverge.

Behavioral analyses included participants with complete data in all three conditions. As in the span+2 analyses, PWA and pupillometry required at least 10 usable span-1 trials within each analysis window, and EEG required at least 15 clean span-1 trials.

#### Behavior

Ninety-nine participants contributed complete behavioral data across the three conditions. Mental demand ratings were available for 80 of them. Full sequence accuracy declined from 84.1% at span-1 to 72.0% at span and 51.2% at span+1, whereas subjective mental demand increased from 8.79 to 10.51 and 12.55, respectively (Table S4). ANOVAs showed effects of Condition on full-sequence accuracy, F(1.80, 176.76) = 329.62, p < .001, *η*_p_^2^ = .77; partial accuracy, F(1.66, 162.20) = 198.92, p < .001, *η*_p_^2^ = .67; partial score, F(1.50, 146.67) = 256.75, p < .001, *η*_p_^2^ = .72; and mental demand, F(1.51, 119.10) = 125.98, p < .001, ηp² = .62. Every pairwise comparison was significant for each outcome (Holm-adjusted ps < .001), indicating a behavioral and subjective progression across the levels of individually adjusted difficulty of the task.

**Table S4.** Behavioral accuracy and subjective cognitive load across span-1, span, and span+1 conditions.

| Condition | Mental demand, M (SD) | Partial-recall accuracy, %, M (SD) | Digits correct, n, M (SD) | Median sequence length | Correct sequences, %, M (SD) |
|---|---|---|---|---|---|
| Span-1 | 8.79 (3.75) | 94.1 (4.71) | 5.51 (1.33) | 6 | 84.1 (9.71) |

*Note. N = 99 for the behavioral outcomes and N = 80 for mental-demand ratings.*

#### Pupil

Eighty-four participants met the span-1 pupillometry inclusion criterion. Seventy-six had complete observations for the first-five and tonic last-five analyses, whereas 82 had complete observations for the four-position late phasic analysis. Tonic pupil size in the terminal encoding and retention interval did not differ by Condition (F(1.70, 140.84) = 1.00, p = .360, *η*_p_^2^ = .01; see Figure S4). Across the first five digits, the effects of Condition (F(1.50, 112.66) = 3.54, p = .045, *η*_p_^2^ = .05), Serial position (F(1.53, 114.52) = 71.23, p < .001, *η*_p_^2^ = .49), and their interaction (F(3.02, 226.47) = 5.19, p = .002, *η*_p_^2^ = .07) were significant. The three conditions were similar through the fourth digit; at the fifth digit, tonic pupil size was larger in span-1 than in span (p = .008) and span+1 (p < .001).

For the last five span-1-matched digits, tonic pupil size showed effects of Condition (F(1.49, 112.03) = 10.21, p < .001, *η*_p_^2^ = .12), Serial position (F(1.26, 94.84) = 37.99, p < .001, *η*_p_^2^ = .34), and their interaction (F(2.82, 211.75) = 27.96, p < .001, *η*_p_^2^ = .27). No condition differences were present at the first two aligned positions. Differences emerged thereafter and culminated at the final span-1 digit, where tonic pupil size followed span-1 > span > span+1 (all ps < .001).

Phasic responses did not show a main effect of Condition at the final digit (F(1.76, 145.87) = 2.90, p = .065, *η*_p_^2^ = .03) or across the first five digits (F(1.49, 111.85) = 2.14, p = .136, *η*_p_^2^ = .03), the first-five Serial Position × Condition interaction was also nonsignificant (F(5.85, 438.74) = 1.37, p = .228, *η*_p_^2^ = .02). In the four-position late analysis (positions −5 through −2; N = 82), Condition (F(1.62, 131.17) = 21.35, p < .001, *η*_p_^2^ = .21) and Serial position (F(2.88, 233.48) = 22.59, p < .001, *η*_p_^2^ = .22) were significant, but their interaction was not (F(4.10, 331.82) = 2.04, p = .087, *η*_p_^2^ = .03). Averaged across these four positions, phasic responses followed span-1 > span > span+1 (span-1 vs. span and span+1, ps < .001; span vs. span+1, p = .011; Table S5).

**Table S5.** Pairwise comparisons of phasic pupil responses averaged across the four late span-1-matched positions.

| Contrast | t | p |
|---|---|---|
| Span-1 - Span | 4.48 | < .001 |
| Span-1 - Span+1 | 5.32 | < .001 |
| Span - Span+1 | 2.60 | .0110 |

Note. The analysis includes aligned positions -5 through -2; position -2 is the final span-1 digit.

**Figure S4.**
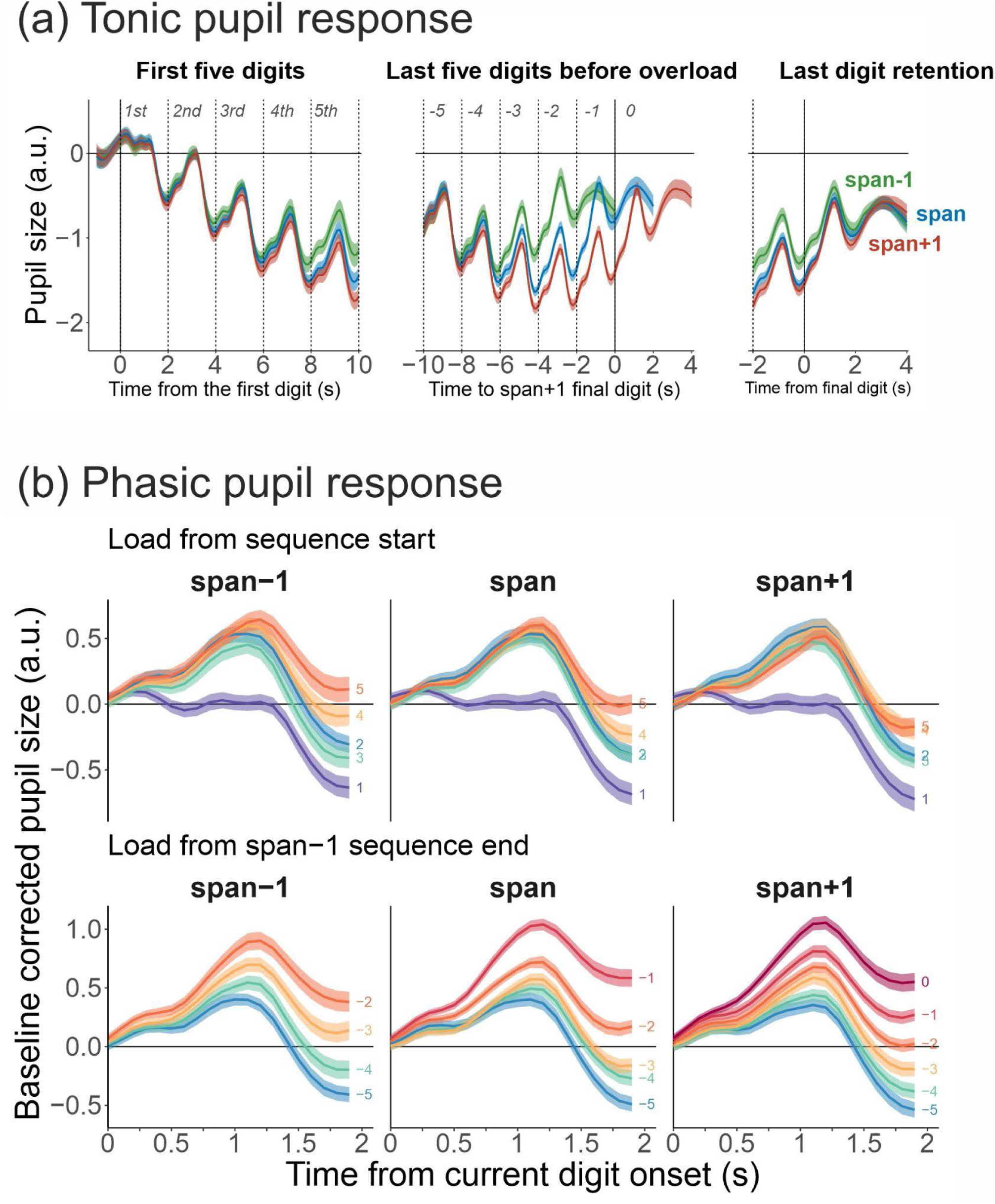
Tonic and phasic pupil responses at span-1, span, span+1. Green denotes span-1, blue denotes span, red denotes span+1. The upper row shows tonic pupil size after subtraction of the 2-s presequence baseline, aligned to sequence onset, to the final span+1 digit, and to each condition’s own final digit. The lower rows show phasic responses baseline-corrected to the 200 ms immediately preceding each digit. Positions 1-5 are counted from sequence onset. Late positions are coded relative to the final span+1 digit: −2 is the final span-1 digit, −1 is the final span digit, 0 is the final span+1 digit. Shaded bands show SEM.

#### PWA

Ninety-three participants contributed complete PWA data for the terminal encoding and retention interval, 89 had complete first- and last-five observations. PWA differed by Condition during terminal encoding and retention (F(1.67, 153.52) = 9.02, p < .001, *η*_p_^2^ = .09; see Figure S5). Span-1 and span did not differ (p = .342), whereas both showed higher PWA than span+1 (span-1 vs. span+1, p = .001; span vs. span+1, p < .001).

Across the first five digits, effects were observed for Condition (F(1.48, 130.64) = 8.68, p = .001, *η*_p_^2^ = .09), Serial position (F(1.63, 143.74) = 23.71, p < .001, *η*_p_^2^ = .21), and their interaction (F(3.28, 288.21) = 4.19, p = .005, *η*_p_^2^ = .05). Span-1 and span did not differ at any position, span differed from span+1 at all five positions, and span-1 differed from span+1 at positions 2-5. In the last-five analysis, Condition (F(1.42, 125.34) = 8.35, p = .002, *η*_p_^2^ = .09) and Serial position (F(1.64, 144.63) = 12.15, p < .001, *η*_p_^2^ = .12) were significant, but their interaction was not (F(3.45, 303.33) = 0.77, p = .530, *η*_p_^2^ = .01). Span-1 and span were similar at four of the five aligned positions, differing only at the earliest position (p = .041), whereas both differed from span+1 throughout (Figure S5). Thus, PWA most closely approximated the predicted within-capacity equivalence (although not formally tested) and overload boundary.

**Figure S5.**
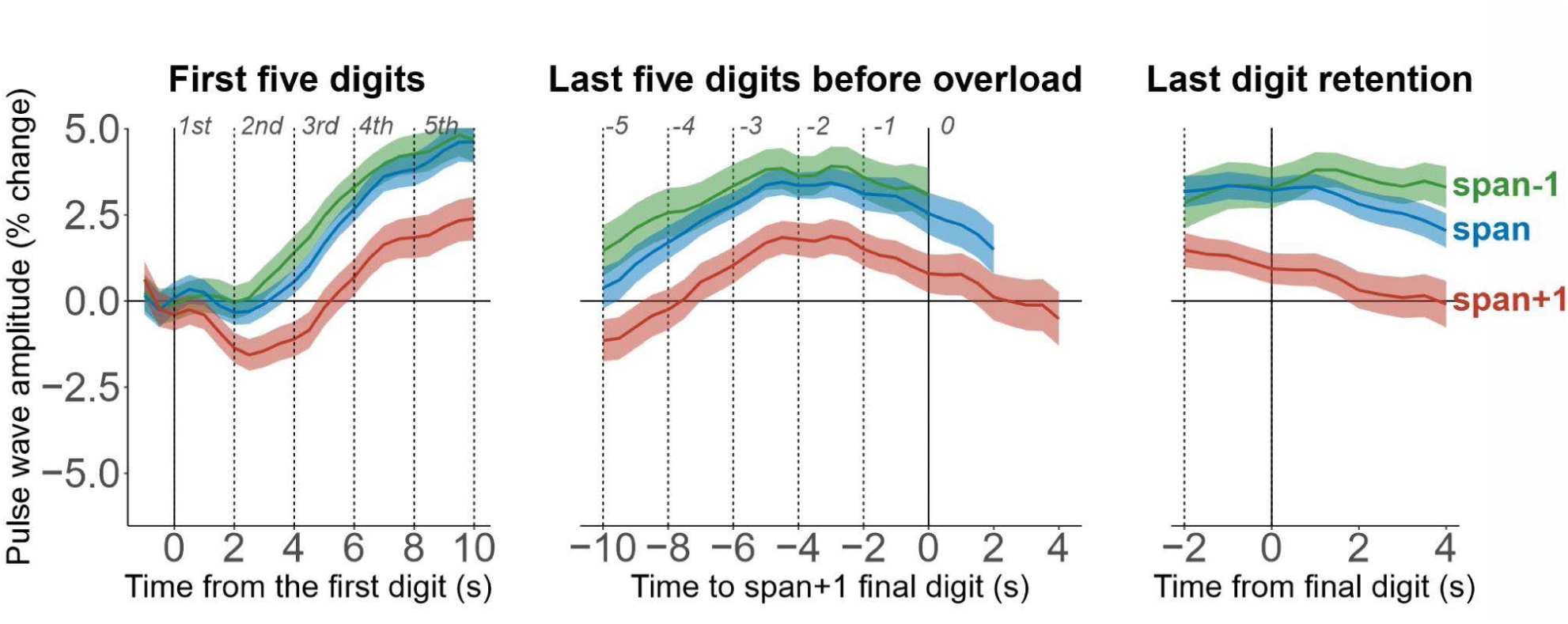
Pulse wave amplitude at span-1, span, span+1. PWA is expressed as percentage change from the 1-s presequence baseline. Green denotes span-1, blue denotes span, red denotes span+1. The left panel is aligned to sequence onset. The middle panel is aligned to the final span+1 digit: −1 is the final span digit, 0 is the final span+1 digit. The right panel realigns each condition to its own final digit. Vertical lines mark digit onsets. Shaded bands show SEM.

#### EEG

Eighty-two participants met the criterion of at least 15 clean span-1 trials. Seventy-six had complete observations for the first five digits, and all 82 had complete observations at the four common late positions (−5 through −2) in each frequency band.

During terminal encoding and retention, frontal theta differed by Condition (F(1.37, 111.32) = 7.27, p = .004, *η*_p_^2^ = .08; see Figure S6): theta power was lower in span-1 than in span (p = .006) and span+1 (p = .005), whereas span and span+1 did not differ (p = .200). Posterior alpha also differed by Condition (F(1.79, 144.65) = 3.46, p = .039, *η*_p_^2^ = .04), with higher power in span-1 than in span (p = .028) and span+1 (p = .038); span and span+1 did not differ (p = .655). Sensorimotor beta showed no terminal condition effect (F(1.53, 124.33) = 0.08, p = .873, *η*_p_^2^ < .01).

Across the first five digits, theta increased and beta became progressively more suppressed as encoding proceeded (main effect of Serial position; see Table S6). These changes followed similar trajectories in all three conditions. Alpha also became progressively more suppressed, but remained higher in span-1 than in span (all ps ≤ .034) and span+1 (all ps ≤ .021) across the five positions. Span and span+1 showed similar alpha power (all ps ≥ .073), and the parallel trajectories indicated that this condition difference was stable across positions.

Across the four common late positions (−5 through −2), theta increased toward the end of encoding in all three conditions (main effect of Serial position; Table S6). Although the increase was numerically smaller in span-1 than in span and span+1, the trajectories did not differ reliably, and there were no pairwise condition differences at any of the four positions (all ps ≥ .182).

At positions −5 through −3, alpha was less suppressed in span-1 than in span (all ps ≤ .023) and span+1 (all ps ≤ .013). Alpha power in span-1 changed little between positions −5 and −4 and then became progressively more suppressed from −4 to −2, whereas it remained comparatively stable in span and span+1. Consequently, the condition means largely converged at position −2, where the span-1 versus span and span-1 versus span+1 comparisons were no longer significant (p = .453 and p = .363, respectively). Span and span+1 remained similar throughout the interval (all ps ≥ .391).

Beta became progressively more suppressed in all conditions, but the decline was steepest in span-1, intermediate in span, and shallowest in span+1. There were no pairwise differences through position −3 (all ps ≥ .060). At position −2, span-1 showed stronger beta suppression than span (p < .001) and span+1 (p < .001), while span showed stronger suppression than span+1 (p = .012; Table S6; Figure S6).

**Table S6.**
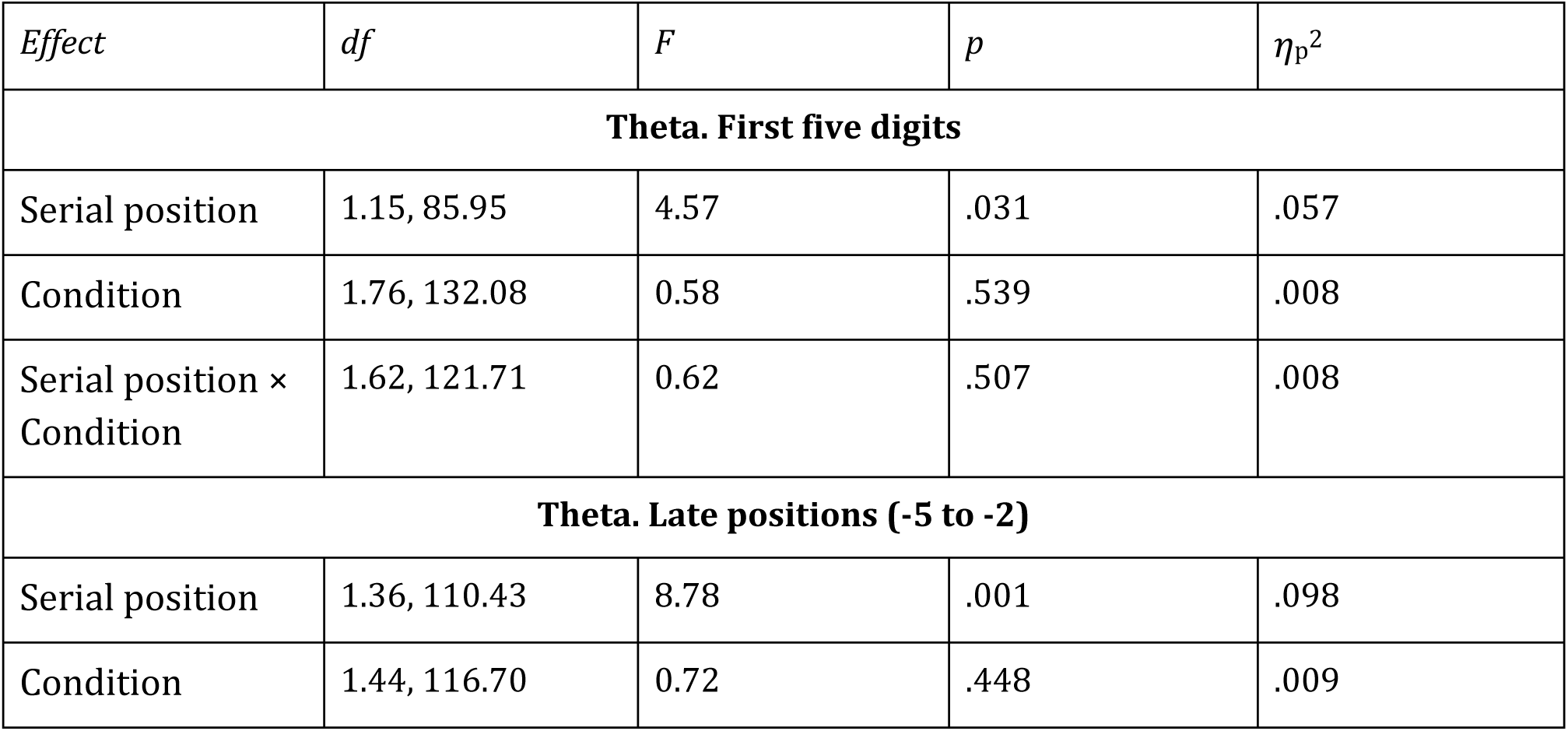
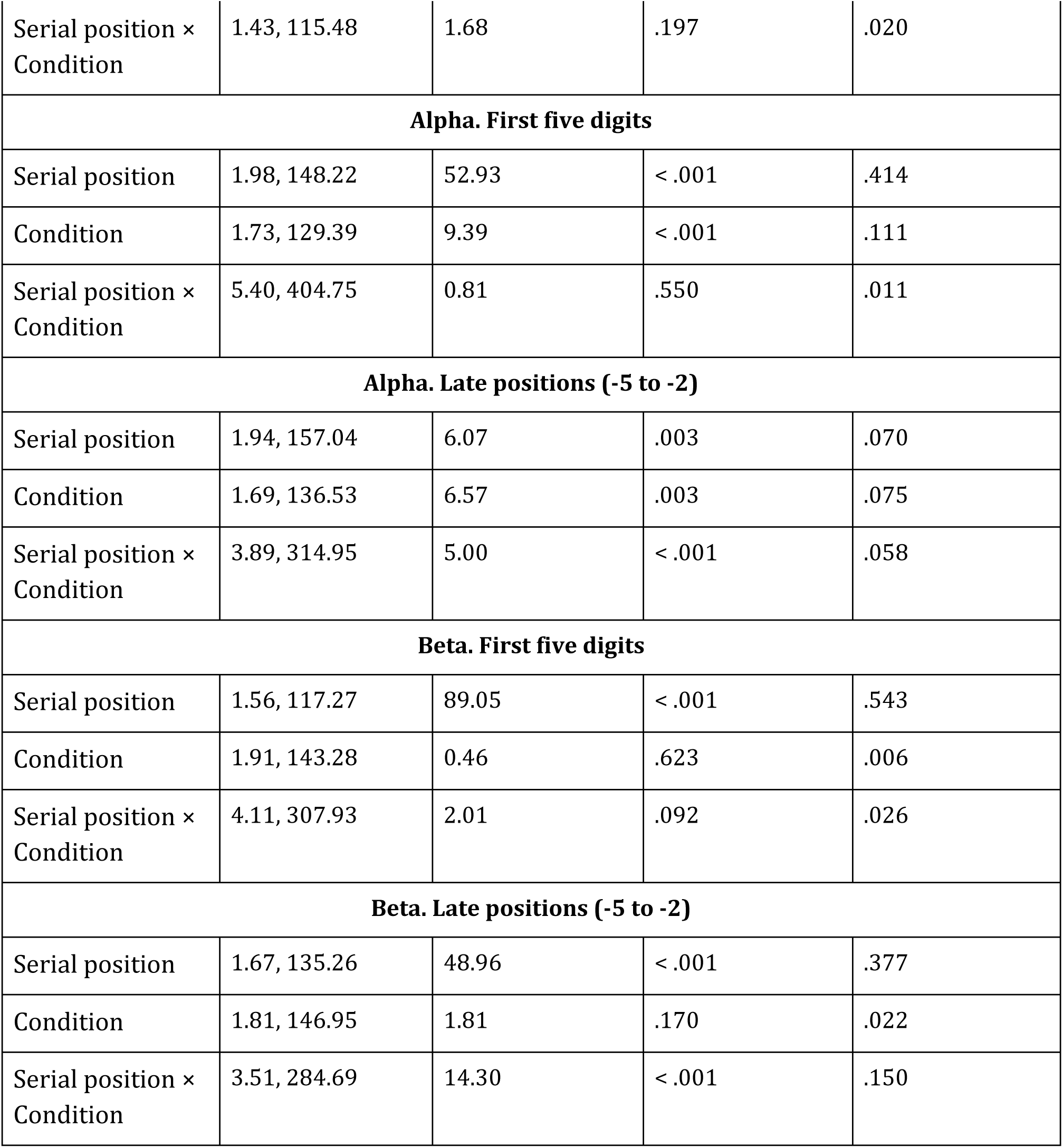
EEG ANOVA statistics for serial position and condition across span-1, span, and span+1 trials.

| *Effect* | *df* | *F* | *p* | *η*<sub>p</sub><sup>2</sup> |
|---|---|---|---|---|
| **Theta. First five digits** |  |  |  |  |
| Serial position | 1.15, 85.95 | 4.57 | .031 | .057 |
| Condition | 1.76, 132.08 | 0.58 | .539 | .008 |
| Serial position × Condition | 1.62, 121.71 | 0.62 | .507 | .008 |
| **Theta. Late positions (-5 to -2)** |  |  |  |  |
| Serial position | 1.36, 110.43 | 8.78 | .001 | .098 |
| Condition | 1.44, 116.70 | 0.72 | .448 | .009 |
| Serial position × Condition | 1.43, 115.48 | 1.68 | .197 | .020 |
| **Alpha. First five digits** |  |  |  |  |
| Serial position | 1.98, 148.22 | 52.93 | < .001 | .414 |
| Condition | 1.73, 129.39 | 9.39 | < .001 | .111 |
| Serial position × Condition | 5.40, 404.75 | 0.81 | .550 | .011 |
| **Alpha. Late positions (-5 to -2)** |  |  |  |  |
| Serial position | 1.94, 157.04 | 6.07 | .003 | .070 |
| Condition | 1.69, 136.53 | 6.57 | .003 | .075 |
| Serial position × Condition | 3.89, 314.95 | 5.00 | < .001 | .058 |
| **Beta. First five digits** |  |  |  |  |
| Serial position | 1.56, 117.27 | 89.05 | < .001 | .543 |
| Condition | 1.91, 143.28 | 0.46 | .623 | .006 |
| Serial position × Condition | 4.11, 307.93 | 2.01 | .092 | .026 |
| **Beta. Late positions (-5 to -2)** |  |  |  |  |
| Serial position | 1.67, 135.26 | 48.96 | < .001 | .377 |
| Condition | 1.81, 146.95 | 1.81 | .170 | .022 |
| Serial position × Condition | 3.51, 284.69 | 14.30 | < .001 | .150 |

**Figure S6.**
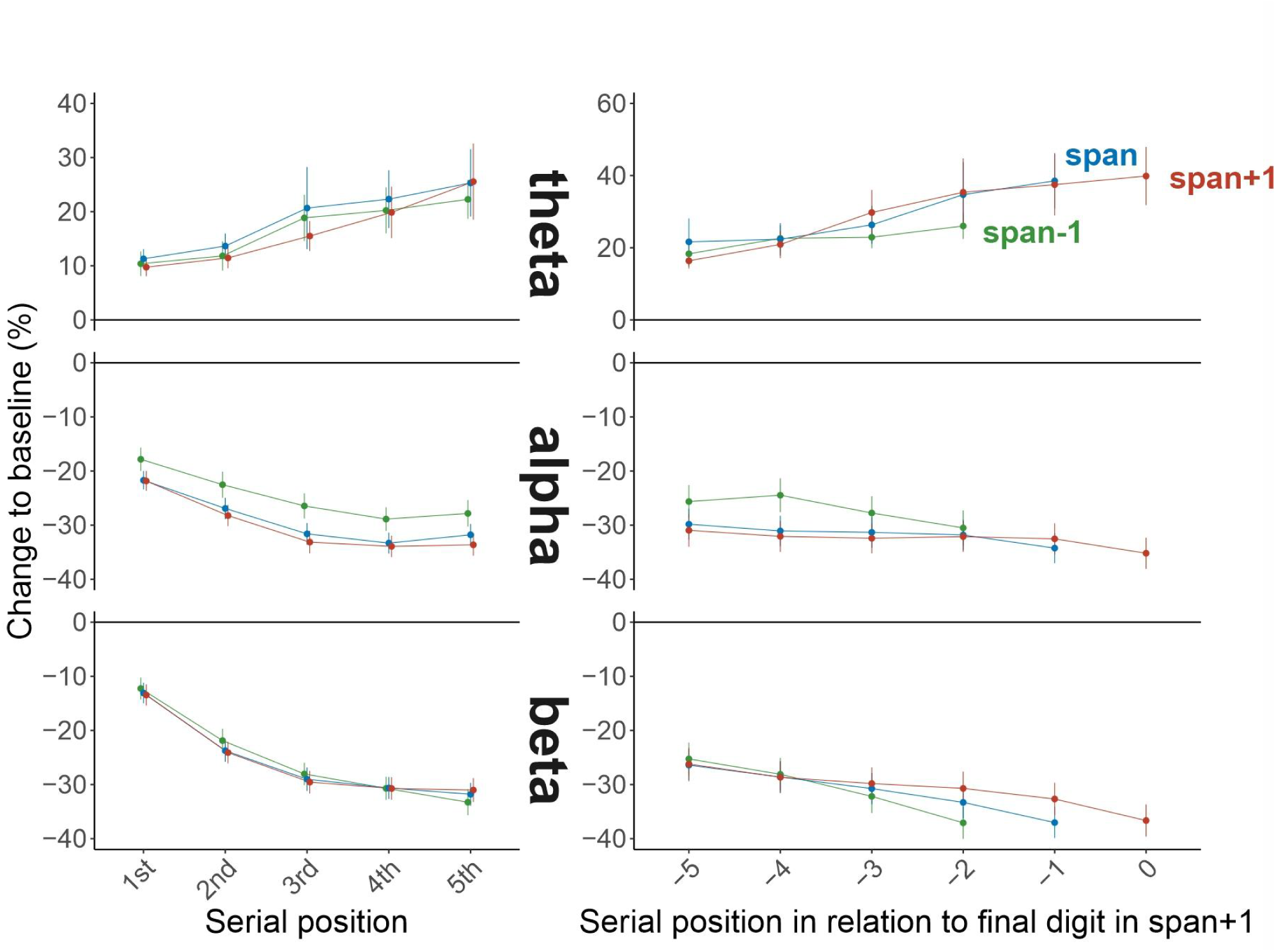
Theta, alpha, and beta power across span-1, span, span+1. Group means are shown for frontal midline theta, posterior alpha, and left centroparietal beta power, expressed as percentage change from the presequence baseline. Green denotes span-1, blue denotes span, red denotes span+1. The left column shows positions 1-5 from sequence onset. In the right column, positions are coded relative to the final span+1 digit: −1 is the final span digit, 0 is the final span+1 digit. Error bars show SEM.

In summary, behavior and subjective demand changed monotonically from span-1 through span+1. PWA provided the clearest evidence for a physiological boundary: span-1 and span were largely similar, while span+1 diverged, although one early aligned late-sequence contrast also separated span-1 from span. Pupillometry showed a more gradual progression across span-1, span, and span+1, with differences emerging mainly during later encoding. EEG followed a different pattern: theta and alpha generally changed from span-1 to span, with little or no additional change at span+1, consistent with saturation at the span level, whereas beta showed an approximately linear difficulty-related divergence only toward the end of encoding. The span-1 analyses therefore support a distinct overload-related PWA pattern, while showing that this categorical transition is not expressed uniformly across physiological systems.

## Subsequent memory effects: Predicting success of the recall from all physiological indicators in span+1 condition

**Table S7.**
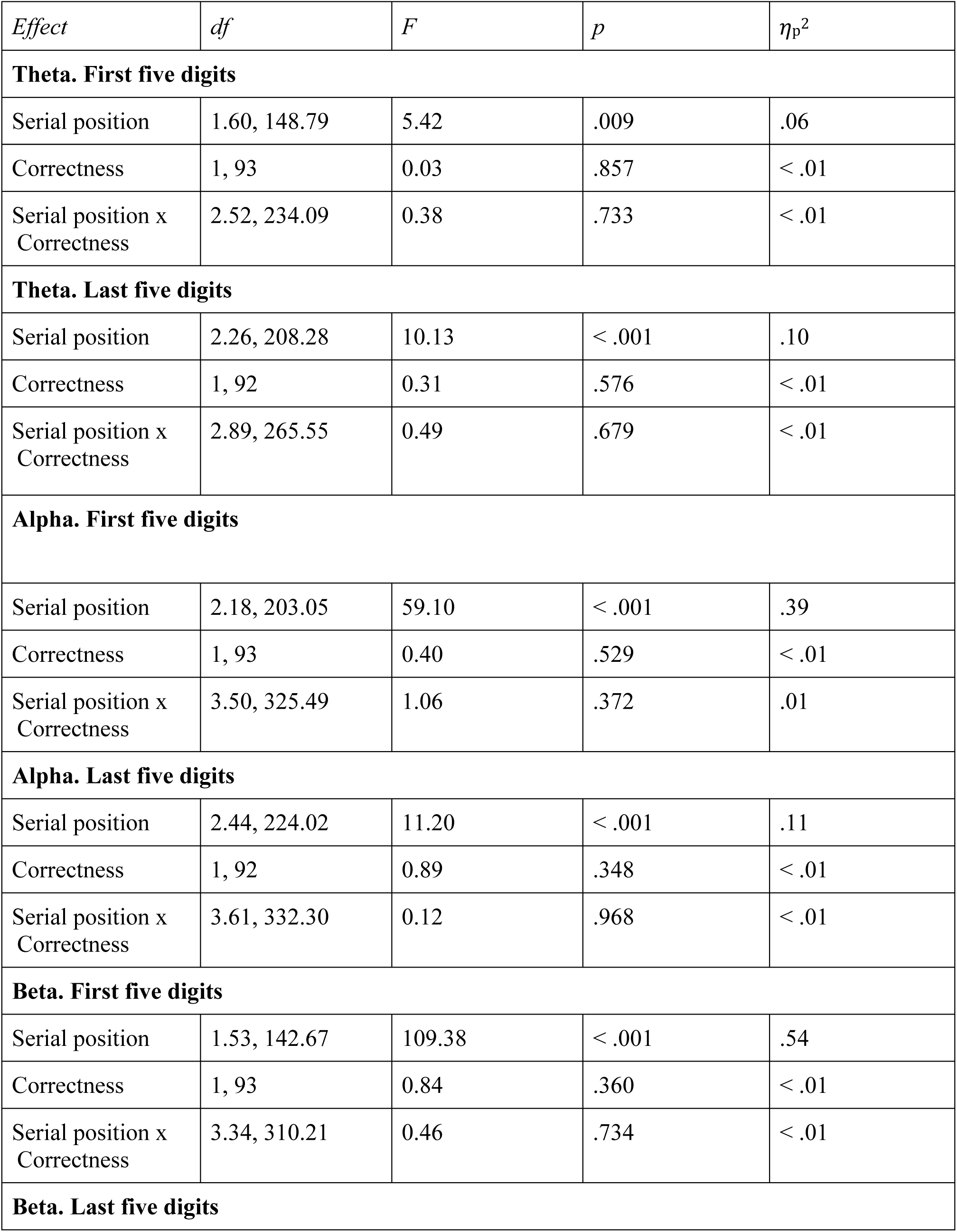
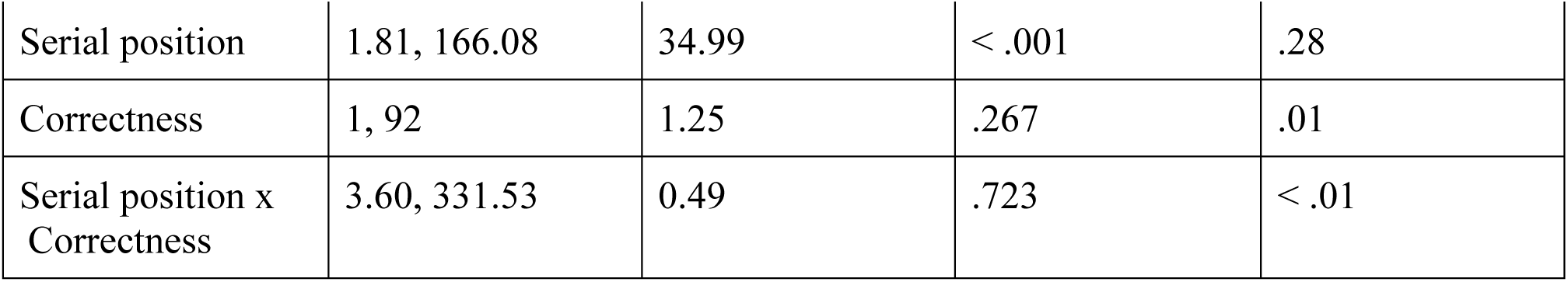
EEG ANOVA statistics for serial position and recall correctness in span+1 trials.

## Notes

### Competing Interest Statement

The authors have declared no competing interest.

